# Left-hemisphere cortical language regions respond equally to observed dialogue and monologue

**DOI:** 10.1101/2023.01.30.526344

**Authors:** Halie Olson, Emily Chen, Kirsten Lydic, Rebecca Saxe

## Abstract

Much of the language we encounter in our everyday lives comes in the form of conversation, yet the majority of research on the neural basis of language comprehension has used input from only one speaker at a time. 20 adults were scanned while passively observing audiovisual conversations using functional magnetic resonance imaging. In a block-design task, participants watched 20-second videos of puppets speaking either to another puppet (the “dialogue” condition) or directly to the viewer (“monologue”), while the audio was either comprehensible (played forward) or incomprehensible (played backward). Individually functionally-localized left-hemisphere language regions responded more to comprehensible than incomprehensible speech but did not respond differently to dialogue than monologue. In a second task, participants watched videos (1-3 minutes each) of two puppets conversing with each other, in which one puppet was comprehensible while the other’s speech was reversed. All participants saw the same visual input but were randomly assigned which character’s speech was comprehensible. In left-hemisphere cortical language regions, the timecourse of activity was correlated only among participants who heard the same character speaking comprehensibly, despite identical visual input across all participants. For comparison, some individually-localized theory of mind regions and right hemisphere homologues of language regions responded more to dialogue than monologue in the first task, and in the second task, activity in some regions was correlated across all participants regardless of which character was speaking comprehensibly. Together, these results suggest that canonical left-hemisphere cortical language regions are not sensitive to differences between observed dialogue and monologue.

## Introduction

Language is first heard, learned and used in informal conversation. Most research on the neural basis of language comprehension, however, has relied on language from a single speaker as stimuli. From the standpoint of a passive observer comprehending language, dialogue between speakers differs from a single speaker in fundamental ways: unlike monologue speech, dialogue is composed of utterances alternating between speakers with different perspectives, voices, and qualities of speech. Comprehending observed dialogue is therefore inherently different from comprehending monologue, and may be an interesting test case for probing the functions of language regions in the brain.

A consistent set of left hemisphere frontal and temporal regions are involved in processing language (Bates et al., 2001; Binder et al., 1997; Dronkers et al., 2004; Fedorenko et al., 2010, 2011; Friederici, 2011; Friederici & Gierhan, 2013; Price, 2010, 2012), robustly responding to language whether it is spoken (Scott et al., 2017), written (Fedorenko et al., 2010), or signed (MacSweeney et al., 2008; Neville et al., 1998; Richardson et al., 2020). These regions in the canonical left-hemisphere cortical language network are active during both production and comprehension (Hagoort, 2014; Hu et al., 2022; Menenti et al., 2011; Price, 2010), in adults and children (Enge et al., 2020), across a wide range of languages (Malik-Moraleda et al., 2022).

They are also sensitive to features of language like comprehension difficulty (Wehbe et al., 2021) and syntactic complexity (Blank et al., 2016), responding more to higher syntactic and semantic processing demands (Hagoort & Indefrey, 2014).

Since early lesion studies, it has generally been accepted that these canonical left-hemisphere language regions are *necessary* for language (Broca, 1865; Wernicke, 1874), but there have been long-standing debates about the *specificity* of these regions for language processing, and in particular, what their limits and scope are (Fedorenko & Thompson-Schill, 2014; Monti et al., 2012). Initially, whole brain activation mapping suggested that language engaged regions that were also active for a range of other cognitive tasks (Blumstein & Amso, 2013; Gold & Buckner, 2002; Thompson-Schill et al., 1997). When language regions are functionally localized within individuals (Braga et al., 2020; Fedorenko et al., 2010), however, these regions are not engaged by nonlinguistic compositional or cognitively difficult tasks like working memory, math, music, cognitive control, action observation, or imitation (Fedorenko et al., 2011; Pritchett et al., 2018). Even reading and evaluating the meaning of computer code – which shares features with language processing like the recursive combination of components in constrained ways to form a more complex meaning (Fedorenko et al., 2019) – does not recruit cortical language regions (Ivanova et al., 2020; Liu et al., 2020), providing further evidence that language regions are highly specific to language processing.

Observing and comprehending dialogue is another interesting boundary case for probing the functions of language regions. Compared to monologues or single-source texts, language in turn-taking dialogue exhibits distinctive features that function to coordinate and monitor the creation of common ground (Clark, 1996; Clark & Schaefer, 1989; Fox Tree, 1999; Fusaroli & Tylén, 2016; Tolins & Fox Tree, 2016). Successive utterances not only convey new meaning, but often show how a prior utterance was understood, facilitating rapid correction (Schegloff et al., 1977). In conversation, speakers quickly volley back and forth – alternating about every 2 seconds with only a 200 ms delay between their utterances on average (Levinson, 2016; Stivers et al., 2009) – establishing referents across speaker boundaries and often finishing each other’s sentences (Clark, 1996; Clark & Schaefer, 1989; Clark & Wilkes-Gibbs, 1986). When observing conversation, adults and even young children can accurately predict turn taking (Casillas & Frank, 2017), and although utterances in dialogue are typically not well-formed grammatical sentences, dialogue is easier to comprehend than monologue from a single speaker (Fox Tree, 1999; Garrod & Pickering, 2004).

Representing and tracking the different perspectives of speakers is integral to understanding dialogue and predicting what might come next. Consider this transcribed excerpt from a two-speaker dialogue without speaker boundaries delineated in the text:

> *Well, you see, I’ve never met him, and so if he comes to the door, how will I know that it’s him? Ah. Oh well, it’s easy. For one thing, we’re exactly alike. You are? Yeah! We’re twins! (Source: https://youtu.be/sS7_-h882Ls)*

As a single linguistic stream, this excerpt – which includes sentence fragments and disfluencies – is hard to understand. Yet, when the utterances are assigned to different speakers, the dialogue becomes easily comprehensible:

> ***Ernie: Well, you see, I’ve never met him, and so if he comes to the door, how will I know that it’s him?***

> *Bert: Ah. Oh well, it’s easy. For one thing, we’re exactly alike.*

> ***Ernie: You are?***

> *Bert: Yeah! We’re twins!*

Knowing that there are multiple speakers – and tracking their alternating perspectives – can impact the interpretation of an utterance and the predictability of the subsequent response. It is therefore plausible that the processes that enable an observer to track the alternating perspectives between interlocutors, which are integral to dialogue comprehension, lie within the scope of canonical language regions.

While the majority of neuroimaging research has focused on language from a single source, some studies have begun examining conversation in the brain (for an excellent review, see (Bögels & Levinson, 2017)). Some prior research, for example, has looked at the neural correlates of comprehension in dialogue when the meaning of an utterance depends on the preceding utterance and contextual information. For example, the utterance “it’s hard to give a good presentation” could be a direct response to the question “how difficult is it to prepare a presentation?” (answer: difficult), or an indirect response to the question “what did you think of my presentation?” (answer: not so great; examples adapted from (Bašnáková et al., 2014)). In the brain, regions including dorsal medial prefrontal cortex (DMPFC), right temporoparietal junction (RTPJ), bilateral inferior frontal gyrus (IFG), and right middle temporal gyrus (MTG) responded more to the same utterance when it was an indirect response than when it was a direct response (Bašnáková et al., 2014; Feng et al., 2017). Another study found that left temporal and frontal regions responded more to indirect than direct replies in question-response pairs (Jang et al., 2013); note that this paper did not control for differences in linguistic features between conditions. Individuals with high communicative skills also showed more activation than individuals with low communicative skills for indirect versus direct responses in dialogue, in regions outside either language or theory of mind network (Bendtz et al., 2022). These results suggest that the processing of implied meaning in indirect responses mostly occurs outside of the core language network. However, this conclusion remains uncertain, as these studies did not use subject-specific functional regions of interest (ss-fROIs) to localize language regions. Activation near IFG might imply modulation of the core language network, or it could reflect activation of nearby ‘multiple demand’ regions that respond to task difficulty (Blank et al., 2014; Fedorenko et al., 2012; Fedorenko & Blank, 2020), especially since indirect replies elicited slower reaction times than the direct replies (Feng et al., 2017). As experimental stimuli, auditory question-response pairs are well controlled, but afford limited opportunity to recognize and resolve differences of perspectives between speakers in context.

In the current study, we test the response of language regions to dialogue by taking a maximal contrast approach: comparing responses to a dyad of alternating speakers (dialogue) versus responses to speech from a single speaker (monologue) using rich, naturalistic, multimodal video stimuli. Both the monologue and the dialogue videos involve rich contexts (e.g., different topics, settings), distinct individuals (e.g., unique characteristics, voices, and mannerisms), and communicative information (e.g., language, gestures). One difference is the intended target-in dialogue, the characters are speaking to each other, whereas in the monologue videos, the characters are addressing the viewer. The ‘directedness’ of speech is a salient cue, even for young children who tend to learn better from child-directed speech (Shneidman et al., 2013; Weisleder & Fernald, 2013). In dialogue, there are also additional features not present in monologue: two individuals – with distinct perspectives, knowledge, goals, and beliefs – interact with each other, cooperating to establish common ground in conversation, building off each other’s responses, and sometimes interrupting each other. Multimodal language comprehension, especially in dialogue, is hypothesized to involve both domain-general and domain-specific mechanisms, which leads to faster processing of multimodal than unimodal language (for review, see (Holler & Levinson, 2019)). While domain-specific language regions in the brain may help with comprehension of multimodal dialogue interactions, if these regions are sensitive to features of dialogue other than linguistic content, then we would expect higher responses in these regions to dialogue than monologue.

In this study, we directly compared activity in adults’ left-hemisphere cortical language regions while they watched naturalistic excerpts of dialogue and monologue (**Experimental Task 1**). We created a block-design task with videos of two characters (from *Sesame Street*) engaging in either a dialogue or two separate monologues, with the audio for each utterance played normally (forward) or temporally reversed (backward). The contrast of forward versus backward speech is a standard manipulation of comprehensibility in auditory language tasks (e.g., (Bedny et al., 2011; Moore-Parks et al., 2010; Olulade et al., 2020)). The key innovation is that we played forward versus backward speech temporally aligned to match naturalistic videos. By comparing responses across the four conditions – forward dialogue, forward monologue, backward dialogue, and backward monologue – we could ask whether there was either a main effect of dialogue (versus monologue), or an interaction between dialogue and language comprehensibility (forward versus backward). We predicted that regions sensitive to dialogue processing should show greatest activity when viewing videos of forward dialogue, compared to both forward monologue (contains language but not social interaction required for dialogue) and backwards dialogue (contains dyadic social interaction but not comprehensible language required for dialogue).

To ensure that any differences (or lack thereof) reflect processing in language regions rather than other nearby cortical regions, we identified subject-specific functional regions of interest (ss-fROIs) for language using a separate auditory language localizer task (Scott et al., 2017). Given the multimodal nature of the stimuli and the range of cognitive processes that dialogue comprehension may tap into (Bögels & Levinson, 2017; Holler & Levinson, 2019; Levinson, 2016), individual functional localization was critical to our approach. Individuals vary in the precise spatial location of functionally-specific regions, and different cognitive functions can often lie next to each other (such as language and executive function; (Blank et al., 2014; Fedorenko et al., 2012; Fedorenko & Blank, 2020)), meaning that group-level approaches can mistake distinctive processing in neighboring regions as a single region performing multiple distinct functions (Fedorenko et al., 2010; Kanwisher, 2010; Saxe et al., 2006). Functionally-defined regions of interest ensure that responses are extracted specifically from language-selective regions in each individual. Regions of interest were identified within left frontal regions (orbital part of inferior frontal gyrus [IFGorb], inferior frontal gyrus [IFG], and middle frontal gyrus [MFG]) and temporal regions (anterior temporal [AntTemp], posterior temporal [PostTemp], and angular gyrus [AngG]).

As a point of comparison, we also examined individually-localized functional regions for two other plausible sets of regions that may respond differently to dialogue and monologue: theory of mind (ToM) regions and the right hemisphere homologues of language regions. Compared to processing linguistic input from a single speaker, understanding overheard dialogue requires tracking the differences between at least two speakers’ perspectives; thus, understanding dialogue may rely more on theory of mind-our ability to reason about others’ minds-than understanding monologue. ToM tasks engage a network of regions in right and left temporoparietal junction (RTPJ, LTPJ), middle, ventral, and dorsal parts of medial prefrontal cortex (MMPFC, VMPFC, DMPFC), and precuneus (PC) (Dufour et al., 2013; Saxe & Kanwisher, 2003; Saxe & Powell, 2006). Given that speaker alternations in dialogue require integrating information from two individuals with different mental states (for instance, in the example above, Bert and Ernie differed in their knowledge of what Bert’s brother looks like), we hypothesized that ToM regions might respond more to dialogue than monologue.

We also measured responses in the individually-defined right hemisphere homologues of language regions, which were also selected for responding more to comprehensible than incomprehensible speech with the separate auditory localizer task. Right hemisphere damage can make it more difficult for individuals to make inferences from discourse (Beeman, 1993), and prior work has demonstrated the right hemisphere’s preferential involvement in social and contextual aspects of language processing (Friederici, 2011; Frühholz et al., 2012; Ross & Monnot, 2008; Seydell-Greenwald et al., 2020). Thus, it was also possible that right hemisphere homologues of language regions might be sensitive to features of dialogue conveyed by the context of the multimodal clips, such as visible interactions between the puppets. Another possibility was that regions outside those we functionally localized may be specifically involved in processing comprehensible dialogue, such as regions involved in processing social interactions (Isik et al., 2017). To address this possibility, we also performed a whole-brain analysis to look for areas responsive to “comprehensible dialogue” by identifying clusters of voxels that were specifically identified by the interaction between comprehensibility (forward>backward) and dialogue (dialogue>monologue).

In addition to the blocked-design Experimental Task 1, the same participants also watched a second task, which offered a complementary test of language regions’ sensitivity to local linguistic structure of utterances versus the larger social, contextual, and visual structure of dialogue. The second task (**Experimental Task 2**) consisted of longer (1-3 minute) continuous clips of dialogue between two characters. Within each clip, one of the two character’s utterances was reversed for the entire dialogue, such that one character spoke forwards and the other replied backwards (incomprehensibly). If language regions are sensitive only to the local occurrence of comprehensible language, it should be possible to extract higher responses to individual forward utterances within the alternating dialogue.

To directly test the sensitivity of language regions to longer temporal scales of social, contextual, and visual aspects of dialogue, we used inter-subject correlation (ISC) analysis (Hasson et al., 2004). The critical assumption was that the partially intelligible dialogues preserved many features of fully intelligible dialogues. The visual input was the same for all participants, but the auditory input was not: which character spoke in forward vs. backwards speech, in each video, was flipped for half of the participants. Thus, the reciprocal clips were exactly matched in the temporal structure of changing common ground, discourse roles of questions and answers, and the overall topic of conversation, as well as the visual features that distinguish dialogue, such as two puppets looking at each other and making contingent gestures and movements. The timecourses of left hemisphere language regions were compared across participants who heard the matched, versus reciprocal, audio stream along with each clip. If *only* the temporal structure of comprehensible language drove activation in language regions, then only participants who heard the same audio stream should show correlated activity. If the social and visual features of the clip also influenced activity in language regions, then all participants should show correlated activity to the same clip. This design cannot isolate which features (contextual, social, and/or visual) of the dialogues are driving the response. However, if language regions do *not* show correlated activity across the reciprocal versions of the same dialogue clip, then those regions’ responses must not be sensitive to any of the features of dialogue that are preserved across the two versions. As a point of comparison, we also extracted individuals’ responses in right homologues of language regions, ToM regions, and regions identified from Experimental Task 1 as responding to comprehensible dialogue.

In summary, we used two novel fMRI tasks to probe the sensitivity of individually-defined left-hemisphere cortical language regions to distinctive features of multimodal dialogue in complementary ways. Regions that processes language independent of a dialogue context should respond equally strongly to comprehensible speech, and equally weakly to incomprehensible speech, whether presented as a monologue or dialogue (Experimental Task 1). Second, these regions should respond selectively to the comprehensible speech segments in a dialogue that alternates between forward and backwards speech, even within the frequent alternations of dialogue that render some utterances quite short (Experimental Task 2). Finally, the responses to these alternating dialogue stimuli should be driven only by the timing of the comprehensible speech segments, and not by any other features of the dialogue (Experimental Task 2).

## General Methods

### Preregistration

Methods and hypotheses were preregistered on OSF: https://osf.io/n4ur5/ (validation as language localizer) and https://osf.io/kzdpc/ (analyses of conversation processing). There were a few deviations from the initial preregistrations for the methods, detailed in

## Supplementary Materials

### Participants

We scanned 20 adults (age: mean(SD) = 23.85(3.70) years, range 18-30 years) who were fluent speakers of English, right-handed, and had no MRI contraindications. Recruitment was restricted to adults with access to the MIT campus according to Covid-19 policies. The protocol was approved by the MIT Committee on the Use of Humans as Experimental Subjects. Informed consent was provided by all participants. Participants were compensated at a rate of $30/hour for scanning, which is standard for our lab and imaging center.

### fMRI Tasks of Interest

The two fMRI tasks of interest were (1) “*Sesame Street* - Blocked Language” (SS-BlockedLang; **Experimental Task 1**) and (2) “*Sesame Street* - Interleaved Dialogue” (**Experimental Task 2**). Participants completed both tasks in the same visit, though methods and results pertaining to each task are discussed separately in the sections below (after **General Methods**).

### fMRI Localizer Tasks

We used two publicly-available fMRI tasks to functionally localize higher order language regions and theory of mind regions in individual participants. ***(1) Auditory Language Localizer.*** This task was previously validated for identifying high-level language processing regions (Scott et al., 2017). Participants listened to Intact and Degraded 18-second blocks of speech. The Intact condition consisted of audio clips of spoken English (e.g., clips from interviews in which one person is speaking), and the Degraded condition consisted of acoustically degraded versions of these clips that were completely incomprehensible (i.e., garbled noise) but matched for acoustic properties (for more details, see (Scott et al., 2017)). Participants viewed a black dot on a white background during the task while passively listening to the auditory stimuli. 14-second fixation blocks (no sound) were presented after every 4 speech blocks, as well as at the beginning and end of each run (5 fixation blocks per run). Participants completed two runs, each approximately 6 min 6 sec long. Each run consisted of 16 blocks of speech (8 intact, 8 degraded). ***(2) Theory of Mind Localizer.*** This task was previously validated for identifying regions that are involved in ToM and social cognition (Dodell-Feder et al., 2011). Participants read short stories in two conditions: False Beliefs and False Photos. Stories in the False Beliefs condition described scenarios in which a character holds a false belief (e.g., a girl places shoes under the bed, her mom moves them when the girl is at school, and then the girl returns to look for her shoes). Stories in the False Photos condition described outdated photographs and maps (e.g., a photo of a boy was taken when he had long hair, but since then he has gotten a haircut). For more details, see (Dodell-Feder et al., 2011). Each story was displayed in white text on a black screen for 10 seconds, followed by a 4-second true/false question based on the story (which participants responded to via an in-scanner button box), followed by 12 seconds of a blank screen (there was also a 12-second blank screen at the beginning of the run). Each run contained 10 blocks. Participants completed two runs, each approximately 4 min 40 sec long. Task performance is reported in **Supplementary Materials**.

### Experimental Protocol

Data were acquired from a 3-Tesla Siemens Magnetom Prisma scanner located at the Athinoula A. Martinos Imaging Center at MIT using a 32-channel head coil. The scanning session lasted approximately 90 minutes and included an anatomical scan and 10 functional scans: 4 runs of SS-BlockedLang (**Experimental Task 1**), 2 runs of SS-IntDialog (**Experimental Task 2**), 2 runs of the auditory language localizer (Scott et al., 2017), and 2 runs of the ToM localizer (Dodell-Feder et al., 2011). T1-weighted structural images were acquired in 176 interleaved sagittal slices with 1.0mm isotropic voxels (MPRAGE; TA=5:53; TR=2530.0 ms; FOV=256 mm; GRAPPA parallel imaging, acceleration factor PE = 2). Functional data were acquired with an EPI sequence sensitive to Blood Oxygenation Level Dependent (BOLD) contrast in 3 mm isotropic voxels in 46 interleaved near-axial slices covering the whole brain (EPI factor=70; TR=2000 ms; TE=30.0 ms; flip angle=90 degrees; FOV=210 mm). 185 volumes were acquired per run for SS-BlockedLang (TA=6:18), 262 volumes were acquired per run for SS-IntDialog (TA=8:52), 179 volumes were acquired per run for the auditory language localizer (TA=6:06), and 136 volumes were acquired per run for the ToM localizer (TA=4:40). fMRI tasks were run from a MacBook Pro laptop and projected onto a 16”x12” screen. Participants viewed the stimuli through a mirror attached to the head coil. Isocenter to screen + mirror to eye was 42" + 6" for both eyes. The SS-BlockedLang and SS-IntDialog tasks were run through PsychoPy3 software version 3.2.4. The auditory language localizer and ToM localizer tasks were run through MATLAB version R2019a and PsychToolbox version 3.0.17.

### fMRI Preprocessing and Statistical Modeling

FMRI data were first preprocessed using fMRIPrep 1.2.6 (Esteban et al., 2019), which is based on Nipype 1.1.7 (Gorgolewski et al., 2011). See **Supplementary Materials** for full preprocessing pipeline details. We then used a lab-specific script that uses Nipype to combine tools from several different software packages for first-level modeling. Event regressors were created for each of the task conditions (Intact and Degraded for the auditory language localizer; False Belief and False Photo for the ToM localizer; see below for details on Experimental Task 1 and Experimental Task 2), and for the response period in the ToM localizer task. Each event regressor was defined as a boxcar convolved with a standard double-gamma HRF, and a high-pass filter (1/210 Hz) was applied to both the data and the model. Artifact detection was performed using Nipype’s RapidART toolbox (an implementation of SPM’s ART toolbox). Individual TRs were marked as outliers if (1) there was more than .4 units of frame displacement, or (2) the average signal intensity of that volume was more than 3 standard deviations away from the mean average signal intensity. We included one regressor per outlier volume. In addition, we included a summary movement regressor (framewise displacement) and 6 anatomical CompCor regressors (Behzadi et al., 2007) to control for the average signal in white matter and CSF. We applied a 6mm smoothing kernel to preprocessed BOLD images. The first-level model was run using FSL’s GLM in MNI space. Subject level modeling was performed with in-lab scripts using Nipype. Specifically, FSL’s fixed effects flow was used to combine runs at the level of individual participants. A subject level model was created for each set of usable runs per contrast for each task (up to 4 runs for SS-BlockedLang, and up to 2 runs for SS-IntDialog, auditory language localizer, and ToM localizer). Runs with more than 20% of timepoints marked as outliers were excluded from analysis (1 run of SS-IntDialog in 1 participant and 1 run of the ToM localizer in another participant were excluded for motion). We also excluded 1 run of SS-BlockedLang and 1 run of SS-IntDialog from a participant who reported falling asleep. Output average magnitudes in each voxel in the second level model were then passed to the group level model. Group modeling used in-lab scripts that implemented FSL’s RANDOMISE to perform a nonparametric one-sample t-test of the con values against 0 (5000 permutations, MNI space, threshold alpha = .05), accounting for familywise error.

### Subject-Specific Functional Individual Region of Interest Analysis

We defined subject-specific functional regions of interest (ss-fROIs) for language as the top 100 voxels activated in an individual, within each of six predefined language search spaces, for the Intact>Degraded contrast using the auditory language localizer task (Fedorenko et al., 2010). The six language search spaces in the left hemisphere included: Left IFGorb, Left IFG, Left MFG, Left AntTemp, Left PostTemp, and Left AngG (similar to (Fedorenko et al., 2010); parcels downloaded from https://evlab.mit.edu/funcloc/). We also looked within the mirror of these search spaces in the right hemisphere (i.e., right hemisphere language homologues), which we refer to as right hemisphere homologues of language regions. We used the same method as above to define ss-fROIs for ToM. In this case, the ToM ss-fROI definition task was the ToM localizer (Dodell-Feder et al., 2011) using the False Belief > False Photo contrast. The predefined ToM search spaces included 7 regions ((Dufour et al., 2013); parcels downloaded from http://saxelab.mit.edu/use-our-theory-mind-group-maps): right and left temporoparietal junction (RTPJ, LTPJ), the precuneus (PC), the dorsal, middle and ventral components of the medial prefrontal cortex (DMPFC, MMPFC and VMPFC), and the right superior temporal sulcus (RSTS). Using the ss-fROIs defined based on the localizer tasks, we then extracted the average magnitude per condition from the SS-BlockedLang task, averaged across all usable runs per participant.

## Experimental Task 1: SS-BlockedLang

### Methods

#### Stimuli Design

Our goal was to create a set of stimuli that allowed us to manipulate both comprehensibility and dialogue versus monologue in a 2×2 block task design (**Figure 1**).

**Figure 1:**
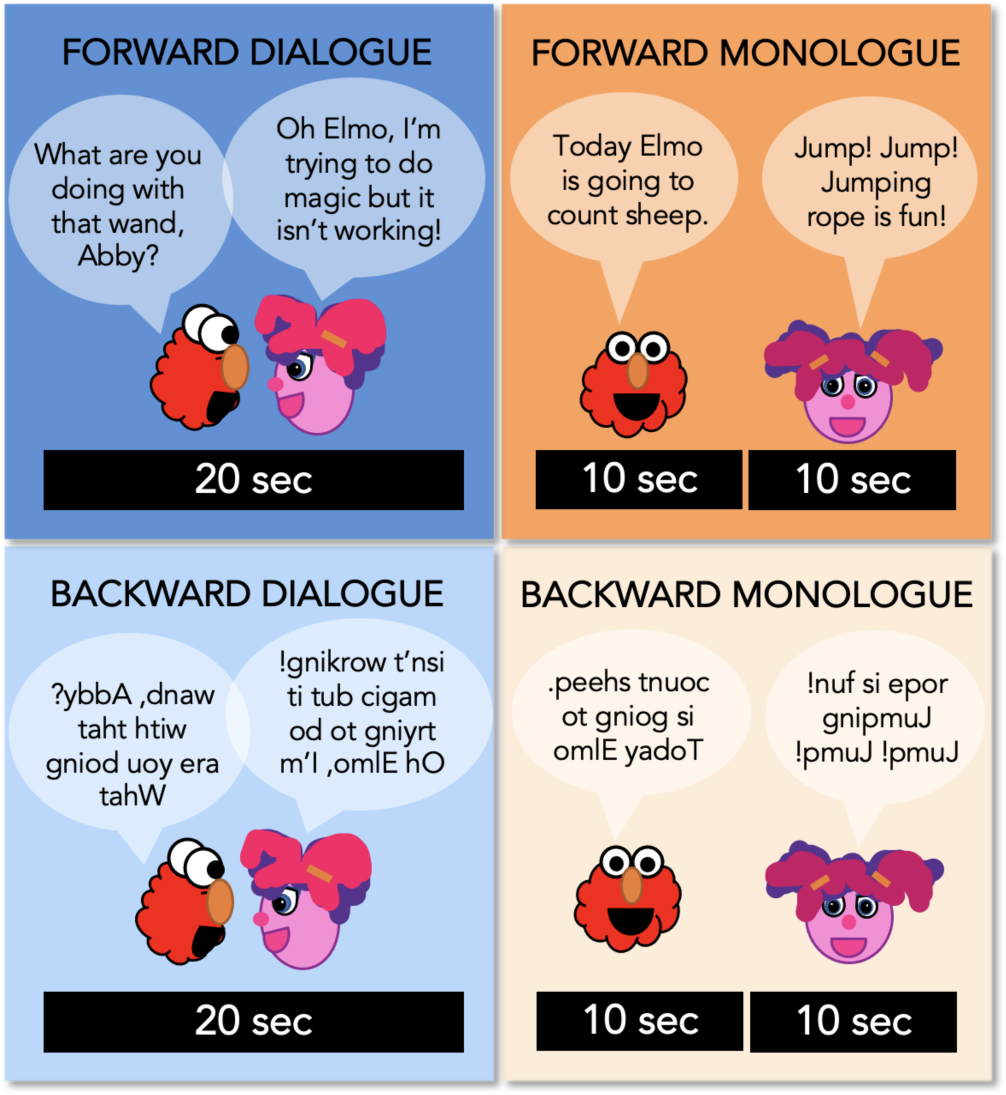
SS-BlockedLang Task Design (Experimental Task 1) Participants watched 20-second clips of Dialogue (blue) and Monologue (orange) of Sesame Street, in which the audio was played either Forward or Backward.

Audiovisual stimuli increase participant engagement with the stimuli, facilitate dialogue comprehension, and emphasize the context of the dialogue by showing two characters interacting on the screen. However, using audiovisual stimuli rather than audio-only stimuli introduced a challenge: how to avoid distracting cross-modal mismatches while varying only the auditory, and not the visual, input across conditions. Even infants and young children are sensitive to the congruence between a speaker’s mouth movements and the sounds they produce in speech (Gogate & Bahrick, 1998; Lewkowicz & Flom, 2014). To balance these desiderata, we used puppets with rigid mouths (rather than human actors) so that the congruence between mouth movements and audio was similar between the forward and backward speech.

We created a set of 20-second edited audiovisual clips of *Sesame Street* during which either two puppets speak to each other (Dialogue), or a single puppet addresses the viewer (Monologue), with the auditory speech stream played either normally (Forward) or reversed so as to be incomprehensible (Backward). Dialogue blocks consisted of two characters, both present in the same scene, speaking back-and-forth for a total of 20 seconds, and Monologue blocks consisted of two sequential 10-second clips of a character, present alone. In the Backward conditions, the audio was reversed within each character rather than across the entire clip, ensuring a continuity of voice-character alignment. For instance, in a Backward Dialogue block with Elmo and Abby, Elmo’s voice was reversed and played when Elmo was talking, and Abby’s voice was reversed and played while Abby was talking.

A notable feature of our task is that it uses commercially produced video clips that were not designed for research purposes. Because we intended to eventually use these same stimuli with very young children, video clips were selected from episodes of *Sesame Street* to appeal to a wide age range. The linguistic content is embedded within colorful, dynamic videos with different characters, different voices, and different settings. To retain the temporal structure and audiovisual match of the clips, the audio was reversed within each utterance of a particular character and carefully overlaid such that the reversed audio still reasonably matched up with the puppets’ mouth movements, and each character’s “voice” was still unique when the audio was reversed. To create the stimuli, we adhered to the following guidelines: (1) we selected only clips that had an overall neutral or positive valence, (2) we included only clips of puppets, rather than clips with humans and puppets, (3) we excluded clips in which the reversed speech did not align well with mouth movements, and (4) we left non-linguistic sounds in the clips, aiming to retain the integrity of the content. We note that there may be residual differences between conditions in the audiovisual alignment that participants may be sensitive to, since the puppets were originally filmed to match the forward speech stream. Transcripts and stimuli features can be found here: https://osf.io/whsb7/

Because we selected commercially available clips, we did not determine the linguistic properties of the stimuli. Monologue and dialogue blocks were matched on the number of mental state words per block, the total number of words per block, and the average age of acquisition for the words per block. However, monologue blocks had significantly longer mean length of utterance, and a lower Flesh-Kincaid reading ease score (see **Supplementary** for details). Notably, even though the dialogue blocks were only 20-seconds long, there were on average more than 6 speaker alternations per block (M(SD)=6.54(2.40), range 2-11).

#### fMRI Task

The SS-BlockedLang task had a 2×2 block design with four conditions: Forward Dialogue, Forward Monologue, Backward Dialogue, and Backward Monologue (**Figure 1**). Participants were asked to watch the videos and press a button on an in-scanner button box when they saw a still image of Elmo appear on the screen after each 20-second block. Participants completed 4 runs, each 6 min 18 sec long. Each run contained unique clips, and participants never saw a Forward and Backward version of the same clip. Each run contained 3 sets of 4 blocks, one of each condition (total of 12 blocks), with each block followed by 1.5 seconds of a still image attention check (Elmo), 0.5 seconds of a blank screen, then either 2 seconds of a fixation cross (within a set of blocks) or 22 seconds of a fixation cross (after each set of 4 blocks; the run also started with a 22-second fixation period). Forward and Backward versions of each clip were counterbalanced between participants (randomly assigned Set A or Set B). Run order was randomized for each participant.

#### Univariate Analysis

For first-level modeling, event regressors were created for each of the four conditions (Forward Monologue, Forward Dialogue, Backward Monologue, Backward Dialogue) and for the button press response period (when a still image of Elmo appeared on the screen and participants were asked to respond via button press). Each event regressor was defined as a boxcar convolved with a standard double-gamma HRF, with the boxcar defined over the onset to the offset of each block. Statistical analyses were conducted in R, using the average activation per condition within ss-fROIs as described in **General Methods**. Conditions were compared using linear mixed effects models; t-tests used Satterthwaite’s method. To test for network-level fixed effects, with ROI and participants modeled as random effects, we used: lmer(mean_topvoxels_extracted∼f_or_b*d_or_m+(1|ROI)+(1|participantID), REML = FALSE), where f_or_b is forwards or backwards (coded 1, −1, respectively), d_or_m is dialogue or monologue (coded 1, −1), and ROI is region of interest within the network. Significance was determined at a level of p<.05 Bonferroni corrected for the three networks tested. To test for interactions within individual regions, we used: lmer(mean_topvoxels_extracted∼f_or_b*d_or_m+(1|participantID), REML = FALSE).

Significance was determined at a level of p<.05 Bonferroni corrected for the number of ROIs (6 for canonical language regions, 6 for right hemisphere language regions, and 7 for ToM regions). In exploratory analyses, we also modeled left and right language regions together and tested for interactions with hemisphere, both at a bilateral language network level and in individual regions, coding for left or right (coded 1,-1).

#### Exploratory Analyses of Conversation Processing

To determine whether brain regions outside the functionally-localized language and ToM regions were specifically responsive to comprehensible dialogue, we performed a whole-brain analysis using the [Forward Dialogue > Forward Monologue] > [Backward Dialogue > Backward Monologue] contrast. Since there were no significant clusters at the preregistered TFCE-corrected (threshold free cluster enhancement) threshold of p<.001, we report exploratory whole-brain results using an uncorrected threshold of p<.001 (two-tailed, 19 degrees of freedom). We then performed exploratory univariate, ss-fROI analyses in conversation regions of interest, i.e., the regions that responded most to comprehensible dialogue in the whole-brain interaction. We created 10mm spheres around the center of gravity (COG) for each significant cluster with at least 10 voxels from the group-level whole-brain analysis. To create ss-fROIs, an in-lab script iteratively used the z-stat image of each 3/4 combined runs (i.e., each ‘fold’) to determine the top 100 voxels for a given subject, ROI, and contrast (in this case, the “comprehensible dialogue” interaction contrast). Critically, this iterative approach ensured that analyzed responses came from independent data that were not used to select an individual’s top-100 voxels. We then used the cope image from the left-out run of a given iteration to extract the betas per condition from these selected top voxels. Statistical analyses were conducted in R. Conditions were compared using linear mixed effects models; t-tests used Satterthwaite’s method. To test for interactions within regions, we used: lmer(mean_topvoxels_extracted∼f_or_b*d_or_m+(1|participantID), REML = FALSE).

## Results

### Univariate response to task conditions in left-hemisphere language regions

The canonical language network, including all six left-hemisphere language regions defined by the independent auditory language localizer (Scott et al., 2017), showed higher responses to both forward speech conditions than both backward speech conditions, as expected (Forward>Backward: Est.=1.05, S.E.=0.05, t-value=19.14, corrected p-value<.001). This pattern held within each individual ss-fROI (**Figure 2**; **Table 1**; corrected p-values<.001 in every region). There was no main effect of Dialogue compared to Monologue in the canonical left-hemisphere language network (Dialogue>Monologue: Est.=0.08, S.E.=0.05, t-value=1.37, corrected p-value=0.52), nor an interaction between comprehensibility and dialogue (Forward>Backward*Dialogue>Monologue: Est.=0.01, S.E.=0.05, t-value=0.22, corrected p-value=1; individual language ss-fROI results in **Figure 2**; **Table 1**; corrected p-values>.1 in every region for dialogue and interaction).

**Figure 2:**
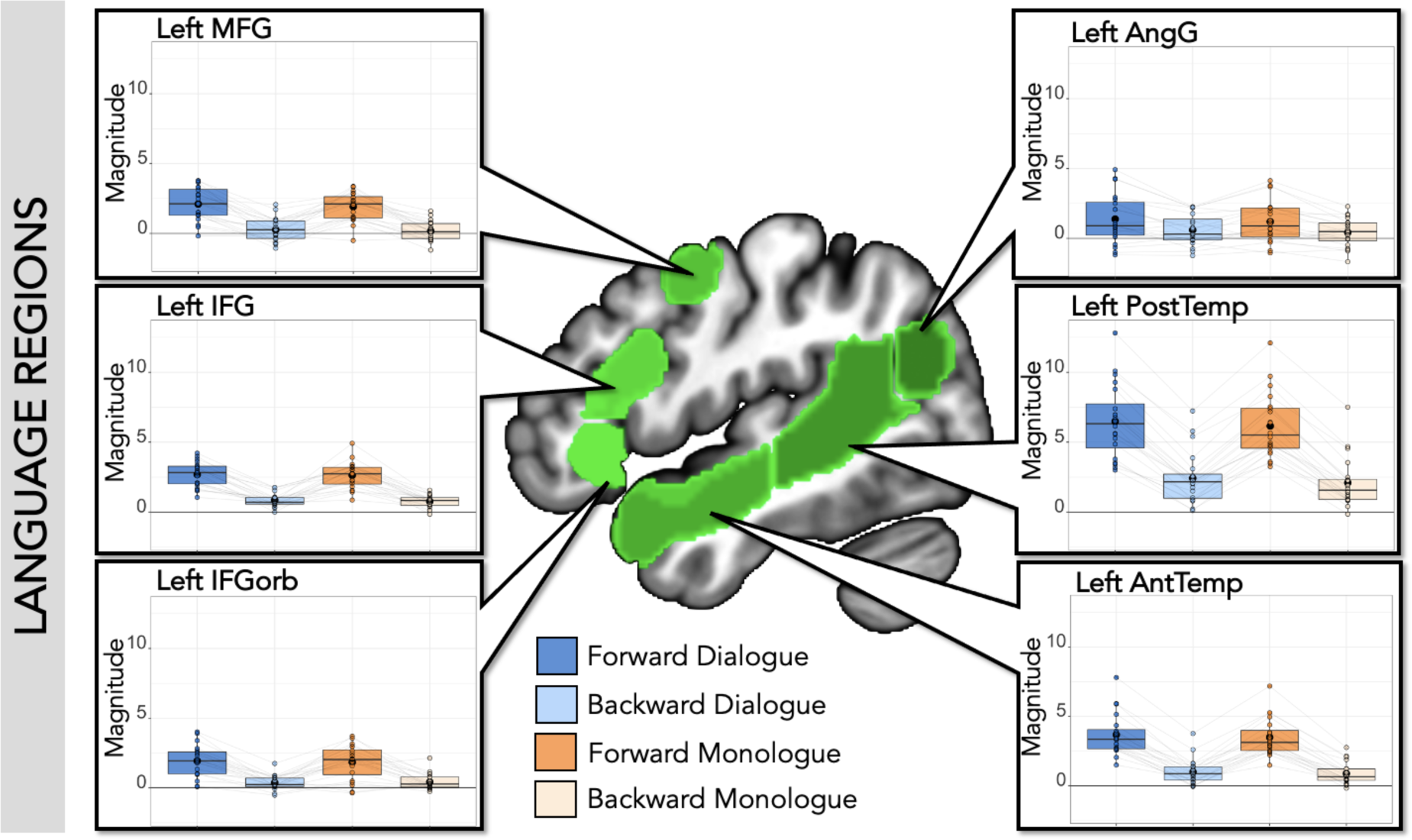
SS-BlockedLang average magnitude by condition within language regions. **Center:** Left hemisphere language parcels overlaid on template brain (green; parcels include left IFGorb, IFG, MFG, AntTemp, PostTemp, and AngG from https://evlab.mit.edu/funcloc/). **Panels**: Average response magnitude (betas) per individual for each condition in the SS-BlockedLang task was extracted from subject-specific functional regions of interest for language (blue: Forward Dialogue; light blue: Backward Dialogue; orange: Forward Monologue; light orange: Backward Monologue). Boxplot with mean in black circle; colored circles show individual participants with light gray lines connecting single participants. There was a main effect of Forward speech compared to Backward speech, but no effect of Dialogue speech compared to Monologue speech within language regions.

**Table 1:**
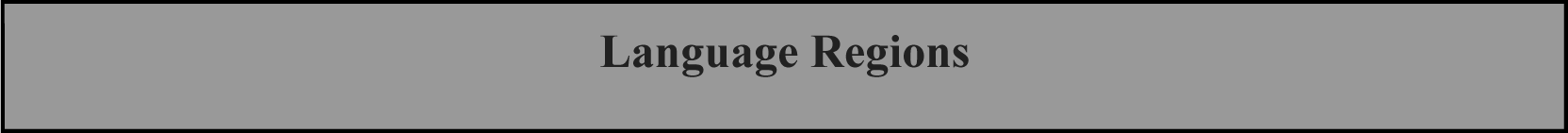
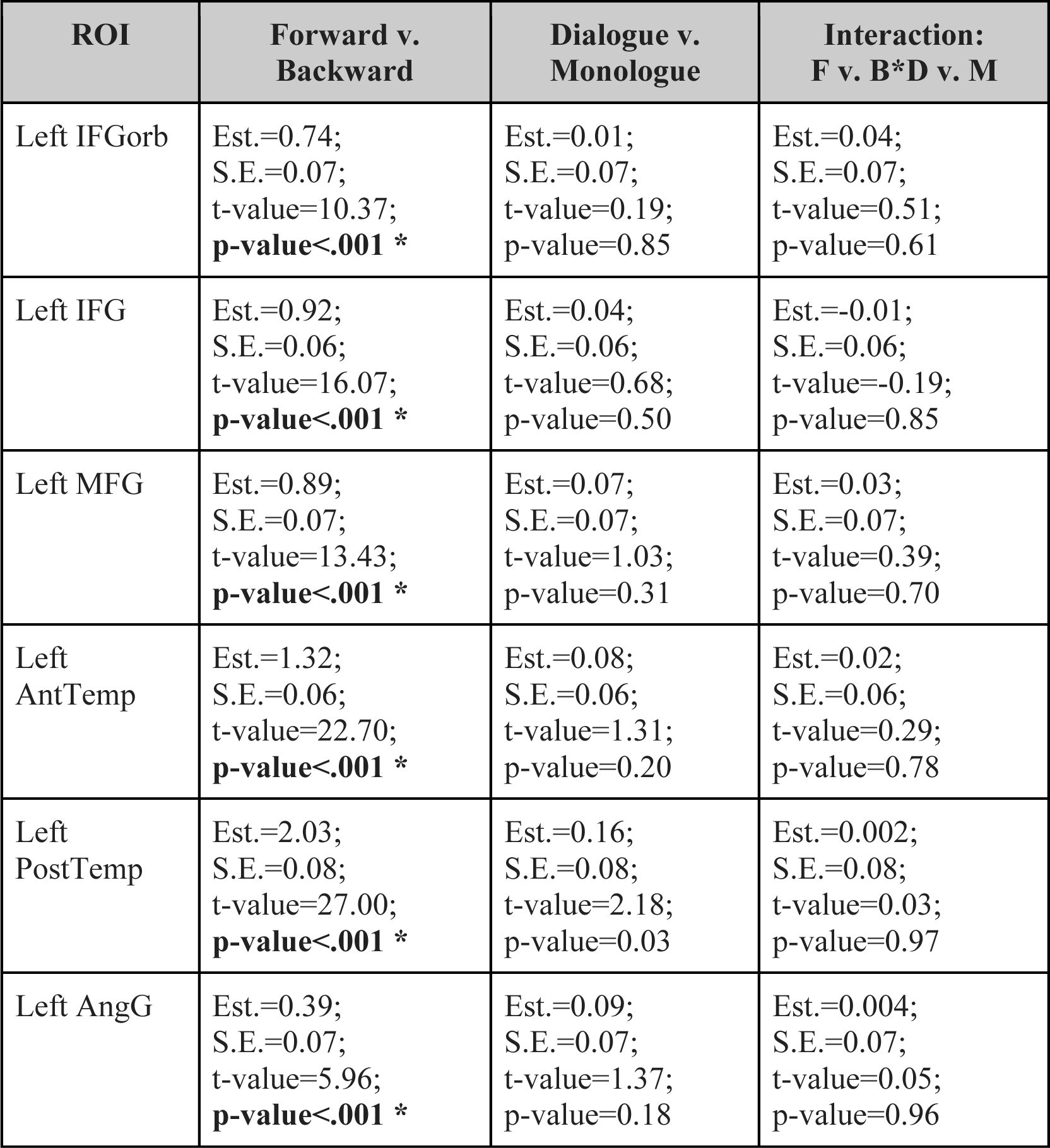
SS-BlockedLang statistics in language regions. Within each language ss-fROI, there was a significant difference between Forward and Backward speech, but no difference between Monologue and Dialogue, and no interaction. Results (Est. = estimate, S.E. = standard error, t-value, and uncorrected p-value) from the model: lmer(mean_topvoxels_extracted∼b_or_f*d_or_m+(1|participantID), REML = FALSE). * indicates significance level p<.05, Bonferroni corrected for 6 ROIs (p<.0083)

### Univariate response to task conditions outside canonical language regions

There were effects of dialogue in regions of cortex outside the canonical left-hemisphere language network. First, we examined right hemisphere homologues of language regions, which responded more to forward than backward speech (Forward>Backward: Est.=0.69, S.E.=0.06, t-value=11.69, corrected p-value<.001), and more to dialogue than monologue speech (Dialogue>Monologue: Est.=0.15, S.E.=0.06, t-value=2.58, corrected p-value=0.03), though showed no interaction between comprehensibility and dialogue (Forward>Backward*Dialogue>Monologue: Est.=0.07, S.E.=0.06, t-value=1.11, corrected p-value=0.80). Individually, all of these regions responded more to forward than backward speech, and right AntTemp and right PostTemp responded more to dialogue than monologue (**Figure 3A**; **Table 2**); there was no significant interaction between comprehensibility (forward/backwards) and dialogue (dialogue/monologue) in any individual region. When both right and left hemisphere language regions were included in the same model, there was a main effect of comprehensibility (Forward>Backward: Est.=0.87, S.E.=0.04, t-value=20.16, uncorrected p-value<.001), a main effect of dialogue (Dialogue>Monologue: Est.=0.11, S.E.=0.04, t-value=2.63, uncorrected p-value=0.01), and an interaction between hemisphere and comprehensibility (Forward>Backward*Left>Right: Est.=0.18, S.E.=0.04, t-value=4.15, uncorrected p-value<.001). For results including hemisphere in the model for individual regions, see **Supplementary Table 1**.

**Figure 3:**
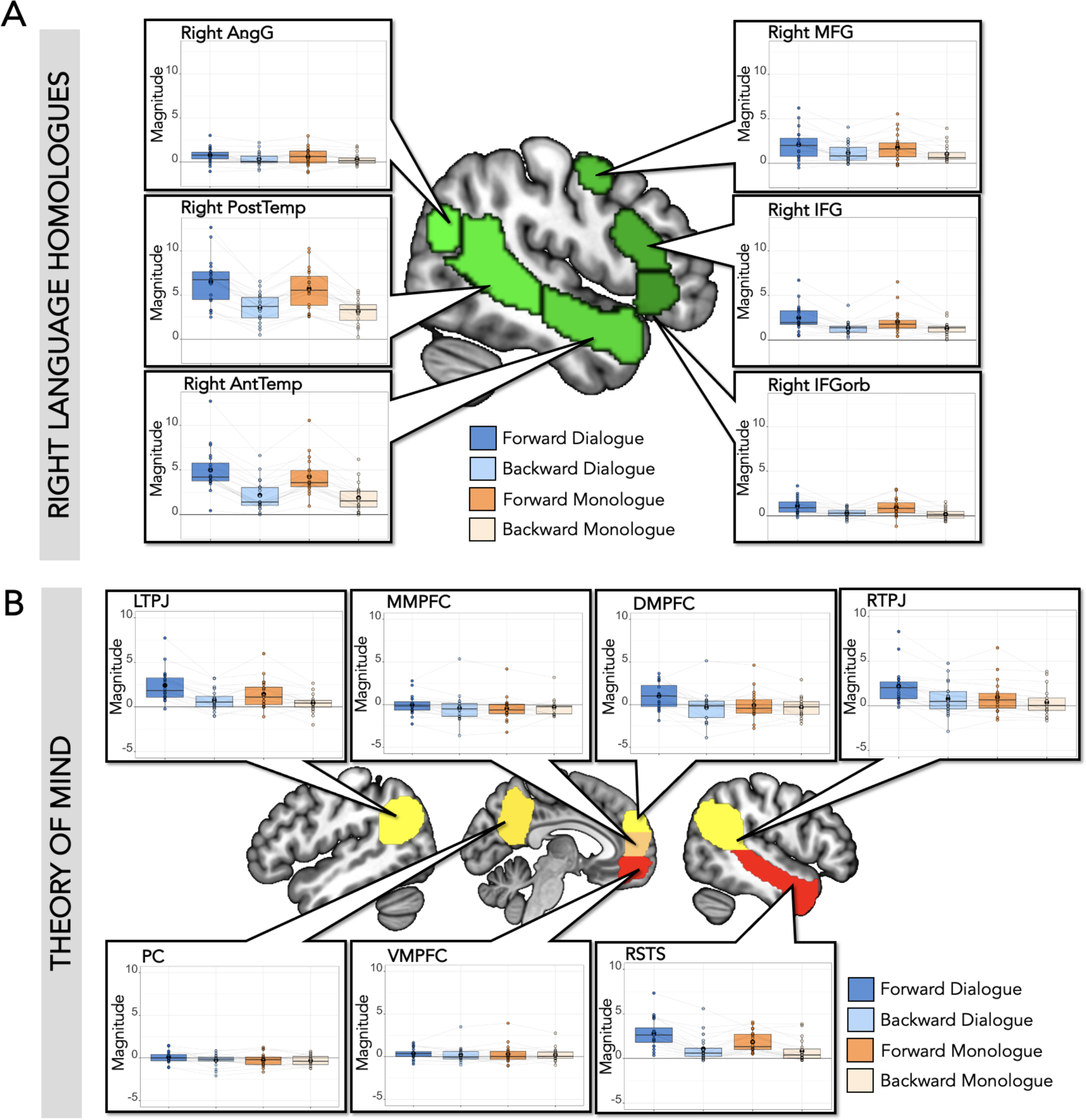
SS-BlockedLang average magnitude by condition within right homologue language regions and ToM regions. **(A) Right Language Homologues. Center**: Right hemisphere language parcels (mirror of left hemisphere parcels) overlaid on template brain (green; parcels include right IFGorb, IFG, MFG, AntTemp, PostTemp, and AngG from https://evlab.mit.edu/funcloc/). **Panels**: Average response magnitude (betas) per individual for each condition in the SS-BlockedLang task was extracted from subject-specific functional regions of interest for right language homologues (blue: Forward Dialogue; light blue: Backward Dialogue; orange: Forward Monologue; light orange: Backward Monologue). Boxplot with mean in black circle; colored circles show individual participants with light gray lines connecting single participants. There was a main effect of Forward speech compared to Backward speech in all regions, and a main effect of Dialogue speech compared to Monologue speech in right AntTemp and PostTemp. **(B) Theory of Mind Regions. Center**: Theory of mind parcels overlaid on template brain (parcels include LTPJ, MMPFC, DMPFC, RTPJ, PC, VMPFC, and RSTS from (Dufour et al., 2013)). **Panels**: Average response magnitude per individual for each condition in the SS-BlockedLang task was extracted from subject-specific functional regions of interest for theory of mind. There was a main effect of Forward compared to Backward speech in DMPFC, LTPJ, RTPJ, and RSTS, a main effect of Dialogue compared to Monologue in LTPJ, RTPJ, and RSTS, and an interaction in DMPFC and RTPJ.

**Table 2:**
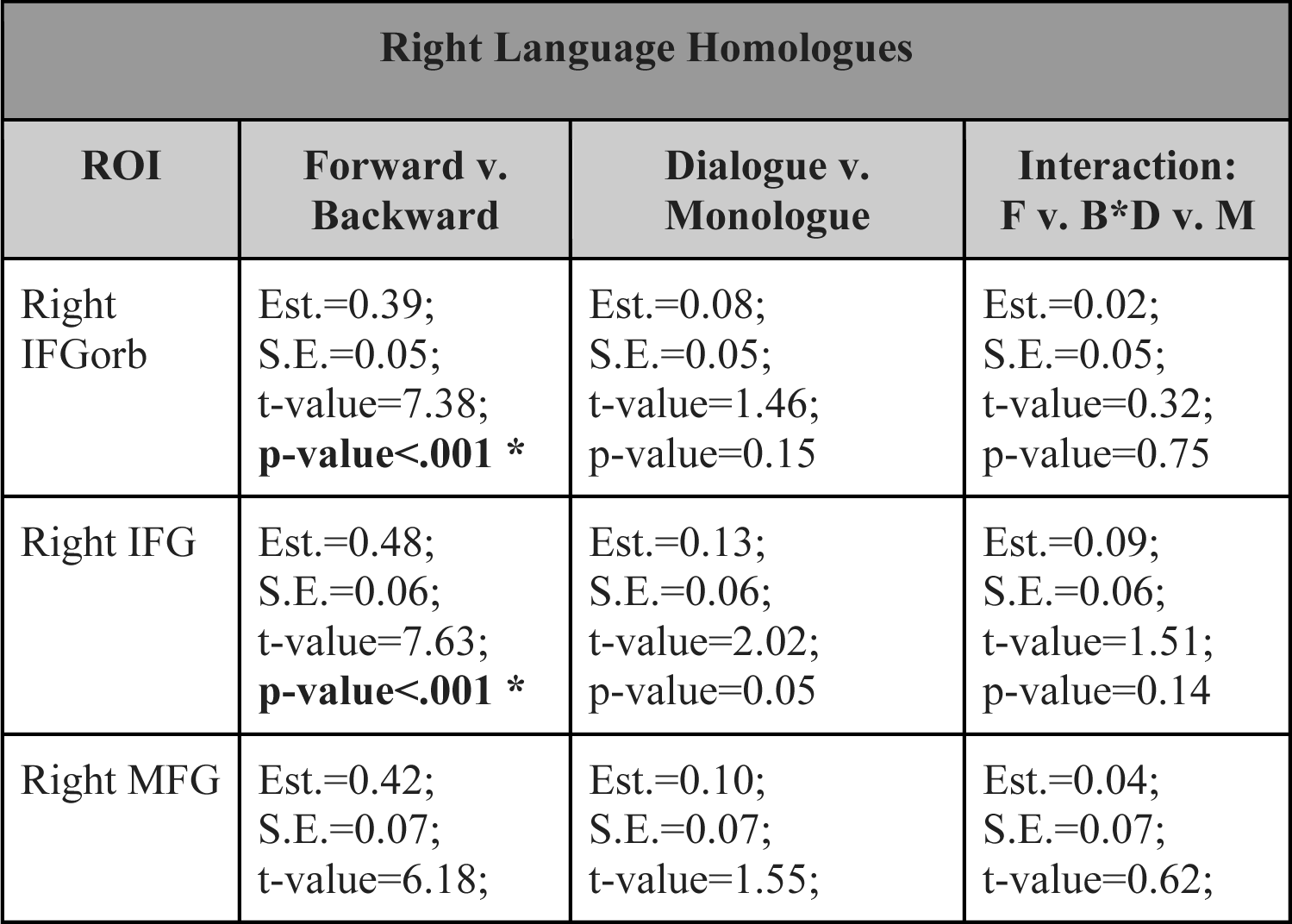
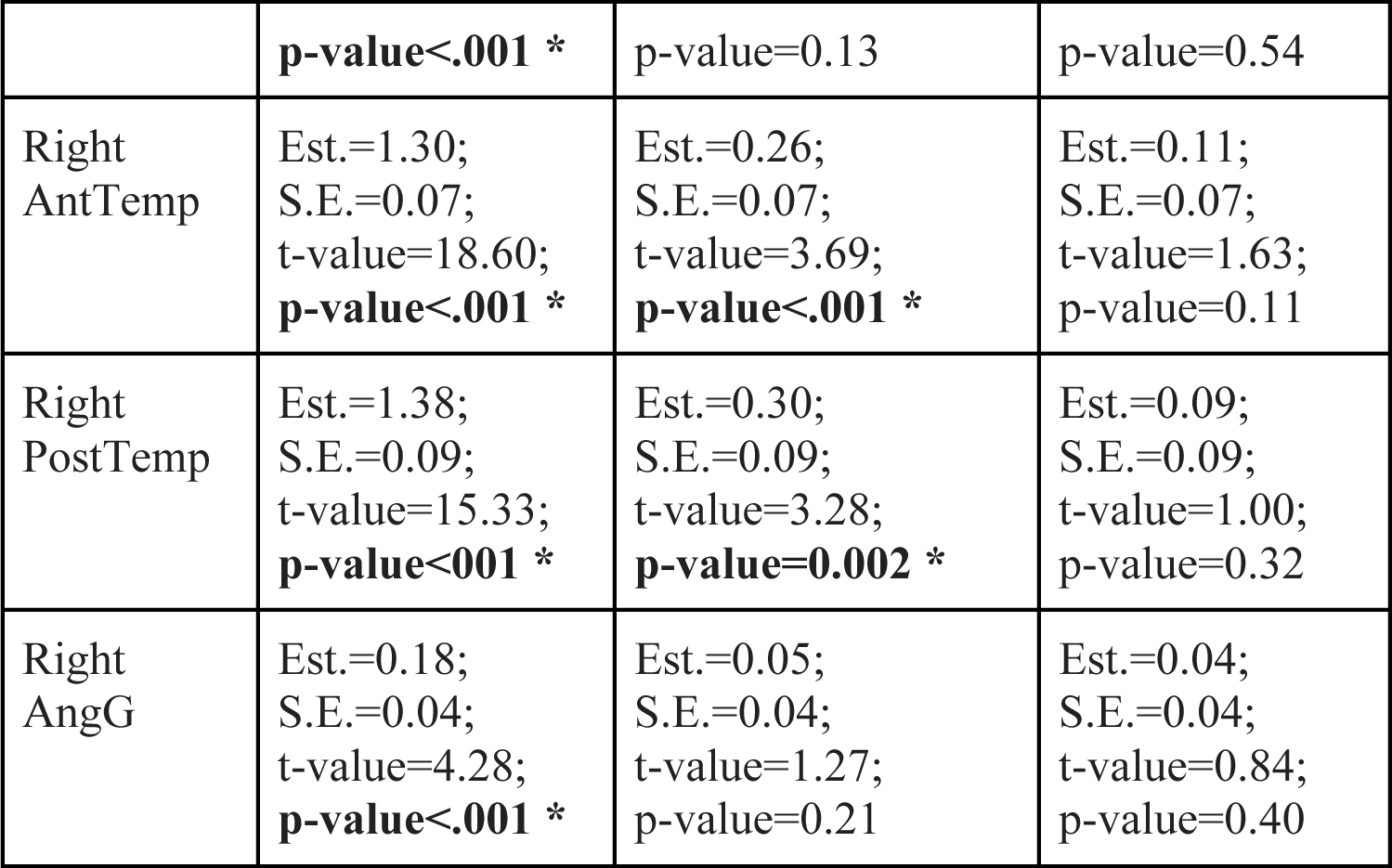
SS-BlockedLang statistics in right hemisphere language region homologues. There was a significant difference between Forward and Backward speech within each right language homologue ss-fROI, and a main effect of Dialogue speech compared to Monologue speech in right AntTemp and PostTemp, but no interaction. Results (Est. = estimate, S.E. = standard error, t-value, and uncorrected p-value) from the model: lmer(mean_topvoxels_extracted∼f_or_b*d_or_m+(1|participantID), REML = FALSE). * indicates significance level p<.05, Bonferroni corrected for 6 ROIs (p<.0083)

Next, we examined responses to each task condition in ToM regions. The ToM network responded more to forward than backward speech (Forward>Backward: Est.=0.35, S.E.=0.05, t-value=6.71, corrected p-value<.001), more to dialogue than monologue (Dialogue>Monologue: Est.=0.21, S.E.=0.05, t-value=4.07, corrected p-value<.001), and showed an interaction between comprehensibility and dialogue (Forward>Backward*Dialogue>Monologue: Est.=0.15, S.E.=0.05, t-value=2.90, corrected p-value=0.01). Individually, four out of seven regions responded more to forward than backward speech (DMPFC, LTPJ, RTPJ, and RSTS), and three responded more to dialogue than monologue (LTPJ, RTPJ, and RSTS; **Figure 3B**; **Table 3**). DMPFC and RTPJ had a significant interaction between comprehensibility and dialogue, responding most to Forward Dialogue. When both left hemisphere language regions and ToM regions were included in the same model, there was a main effect of comprehensibility (Forward>Backward: Est.=0.70, S.E.=0.04, t-value=17.51, p-value<.001), a main effect of dialogue (Dialogue>Monologue: Est.=0.14, S.E.=0.04, t-value=3.59, p-value<.001), a main effect of network (Left_Language>ToM: Est.=0.68, S.E.=0.26, t-value=2.55, p-value=0.02), an interaction between comprehensibility and network (Forward>Backward*Left_Language>ToM: Est.=0.35, S.E.=0.04, t-value=8.74, p-value<.001), and an interaction between comprehensibility and dialogue (Forward>Backward*Dialogue>Monologue: Est.=0.08, S.E.=0.04, t-value=2.05, p-value=0.04).

**Table 3:**
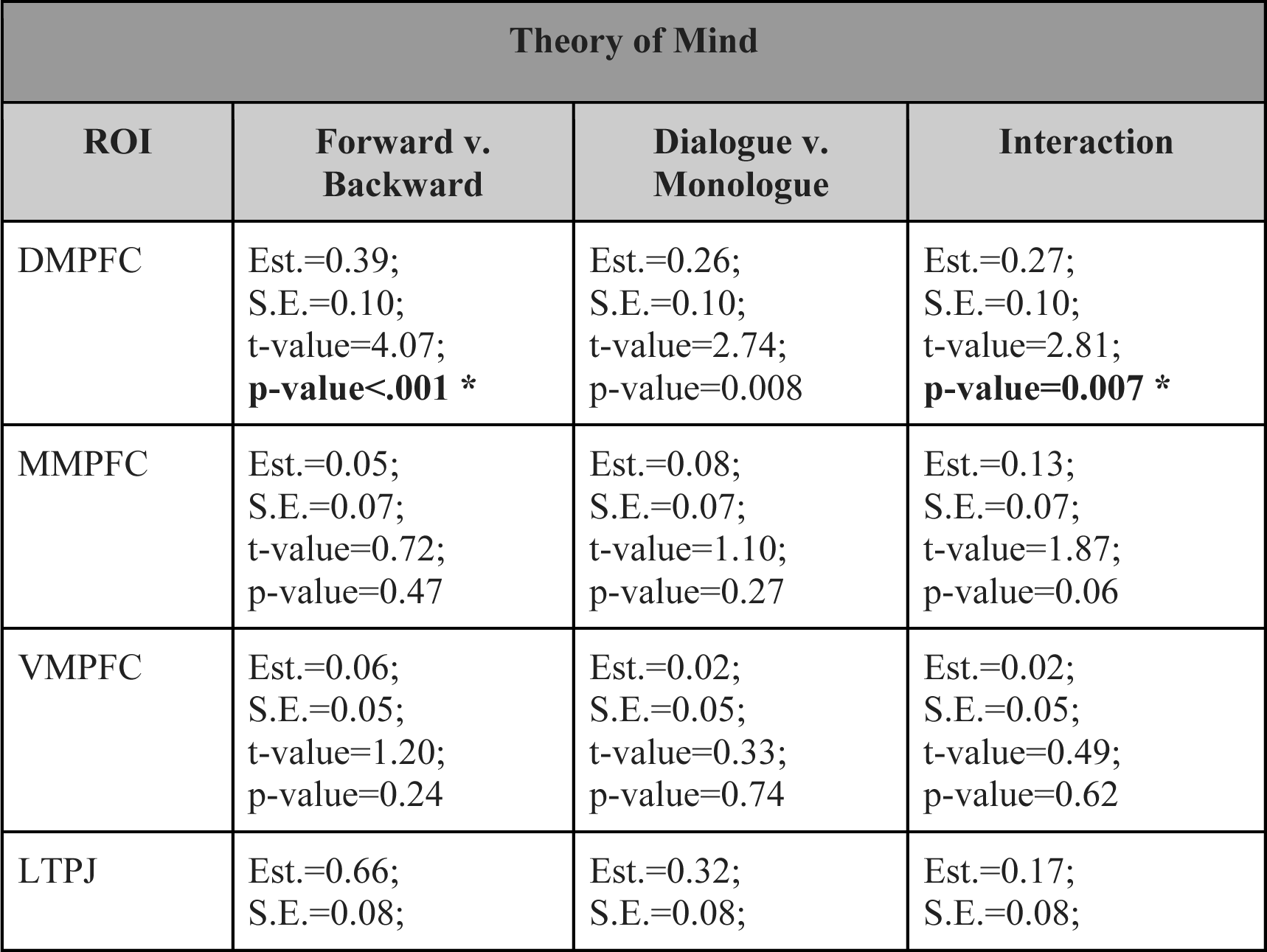
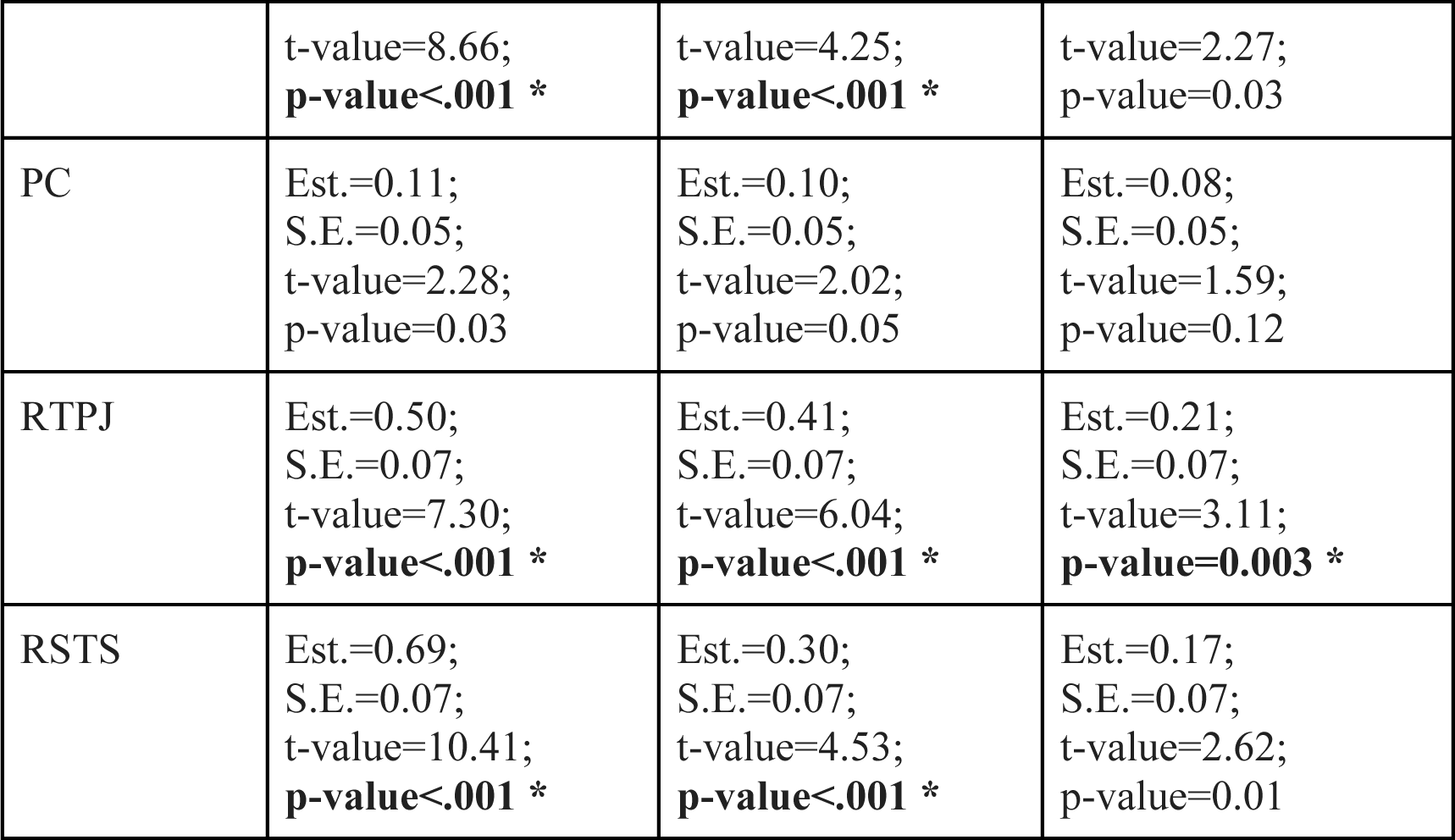
SS-BlockedLang statistics in theory of mind regions. Within ToM ss-fROIs, there was a main effect of Forward compared to Backward speech and a main effect of Dialogue compared to Monologue in DMPFC, LTPJ, RTPJ, and RSTS, and an interaction in DMPFC and RTPJ. Results (Est. = estimate, S.E. = standard error, t-value, and uncorrected p-value) from the model: lmer(mean_topvoxels_extracted∼f_or_b*d_or_m+(1|participantID), REML = FALSE). * indicates significance level p<.05, Bonferroni corrected for 7 ROIs (p<.0071)

Finally, to empirically test for regions that specifically responded to comprehensible dialogue, we performed a whole brain analysis for the following interaction: [Forward Dialogue>Forward Monologue]>[Backward Dialogue>Backward Monologue] (**Figure 4**). Four clusters were identified using an uncorrected threshold of p<.001 in the right temporal pole, right STS, left STS, and left cerebellum (none survived TFCE correction for multiple comparisons). In exploratory analyses, we extracted activity in individual participants in individually defined ss-fROIs (within the 10mm sphere search spaces around center of gravity coordinates from the group results), using a leave-one-run-out approach to maintain independence between data used to define these regions and data used for extracting activation (**Figure 4; Supplementary Table 2**). All four regions responded more to Dialogue than Monologue, and more to Forward than Backward speech. There was an interaction between comprehensibility and dialogue in the right temporal pole and left Crus II (cerebellum).

**Figure 4:**
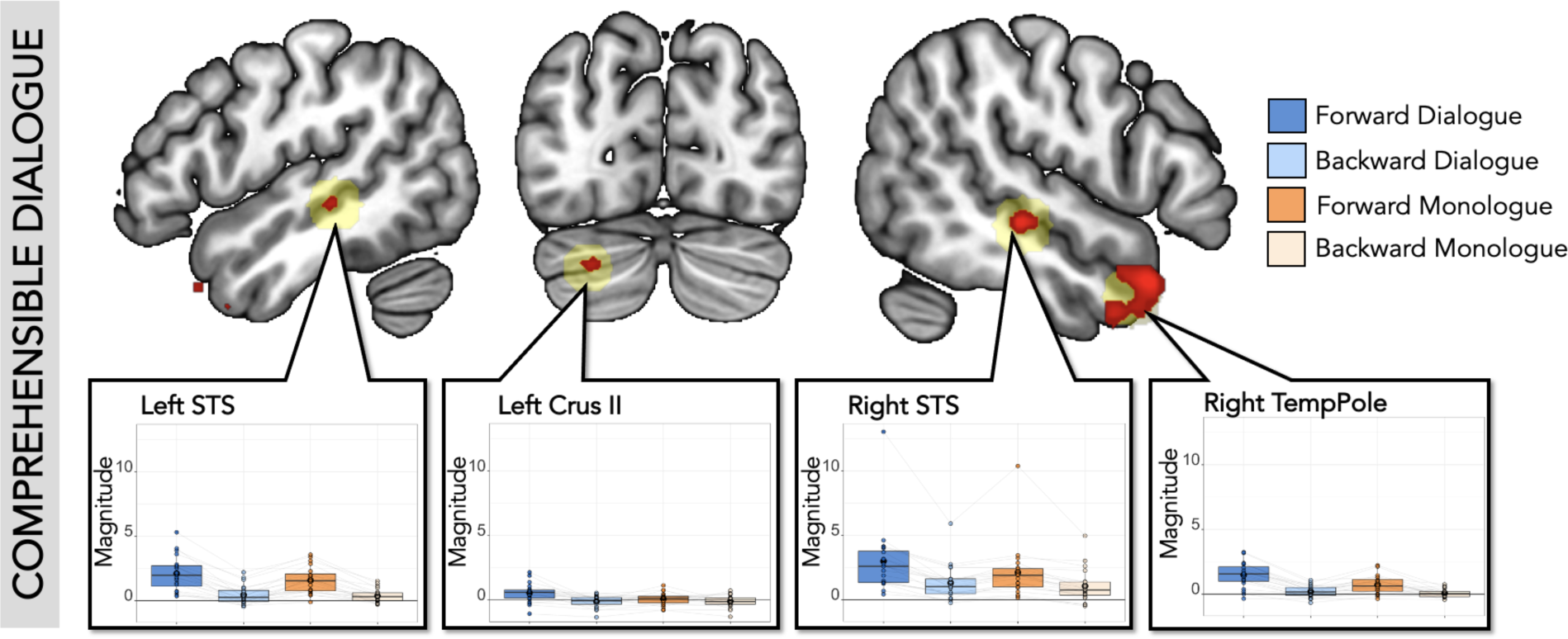
SS-BlockedLang whole brain interaction for comprehensible dialogue. **Top**: In red are significant voxels at a threshold of p<.001, uncorrected (df=19, two-tailed) in the whole-brain analysis for [Forward Dialogue > Forward Monologue] > [Backward Dialogue > Backward Monologue]. We used the uncorrected threshold since nothing survived at TFCE corrected threshold. Significant clusters were identified in right STS, right temporal pole, left STS, and left Crus II (cerebellum). 10mm ROI spheres (light yellow) were created around center of gravity (COG) coordinates from the 4 significant clusters. **Panels**: Average response magnitude per individual for each condition in the SS-BlockedLang task was extracted from comprehensible dialogue ss-fROIs constrained by the spherical ROIs (blue: Forward Dialogue; light blue: Backward Dialogue; orange: Forward Monologue; light orange: Backward Monologue). Boxplot with mean in black circle; colored circles show individual participants with light gray lines connecting single participants. There was a higher response to forward than backward speech in all regions, a higher response to dialogue compared to monologue in all regions, and an interaction in Right Temporal Pole and Left Crus II (cerebellum).

### Summary

These results suggest canonical left-hemisphere cortical language regions do not respond differently to audiovisual dialogues compared to monologues, nor is there an interaction with comprehensibility. The magnitude of response in canonical left-hemisphere language regions appears to be determined only by the presence of comprehensible speech (common to both Forward conditions). In contrast, distinct cortical regions seem to be sensitive to the differences between dialogue and monologue speech, including some ToM regions (DMPFC, LTPJ, RTPJ, and RSTS) and two right-hemisphere homologues of language regions (right AntTemp and PostTemp), as well as other regions identified by exploratory whole-brain analyses (right temporal pole, right STS, left Crus II in cerebellum, and left STS).

## Experimental Task 2: SS-IntDialog

In **Experimental Task 2**, we probed the sensitivity of language regions to features of dialogue by using longer clips of dialogue with interleaved forward and backward speech. Rather than blocks of all-forward and all-backward speech, one character’s audio stream was played forward, while the other character’s audio stream was played backward (which character was forward versus backward was counterbalanced between participants). This approach complements Experimental Task 1. First, we measured canonical language regions’ responses to comprehensible utterances within the temporal structure of natural dialogue, i.e., frequent short utterances, instead of long blocks. Second, and critically, using the inter-subject correlations (ISC), we directly measured the influence of linguistic structure, compared to all other visual and abstract semantic structure of the dialogue, on the timecourse of stimulus-driven activity in canonical language regions.

## Methods

### Stimuli Design

General methods for stimuli design were similar to **Experimental Task 1**. We selected full scenes of dialogue from *Sesame Street* during which two puppets speak to each other (the selected scenes ranged from 1-3 minutes, and we played the entire scene). Like the clips used in **Experimental Task 1** (SS-BlockedLang task), these scenes varied in terms of their visual properties (e.g., objects, setting), topic, and characters. For each clip, we reversed the audio for one character’s utterances, but left the other character’s audio forward (**Figure 5A**).

**Figure 5:**
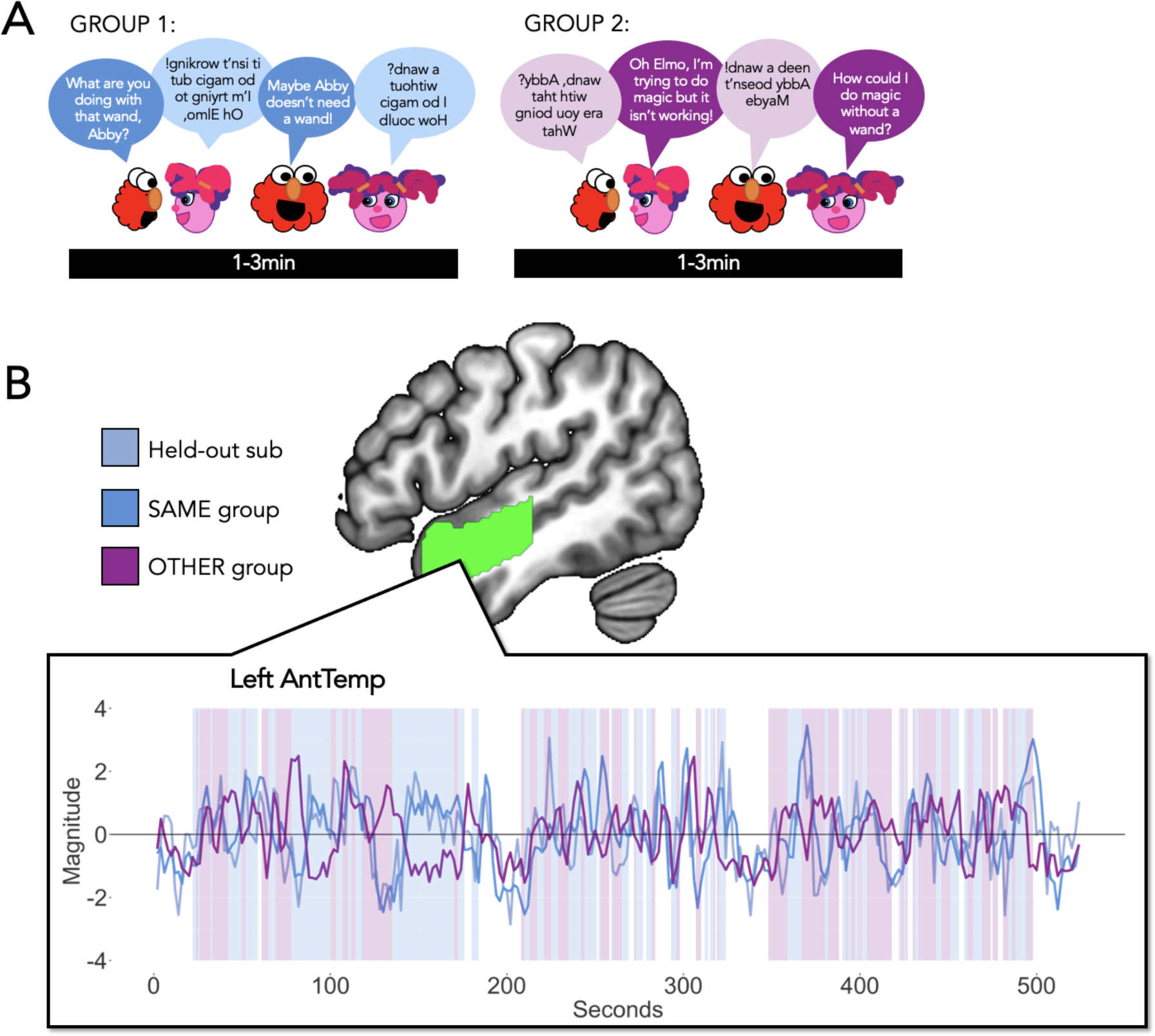
SS-IntDialog Task Design (Experimental Task 2) **(A) Stimuli.** Participants watched 1-3-minute clips of Sesame Street in which two characters have a conversation. The audio from one character was played forward while the second is played backward. Participants were randomly assigned to hear one of the two versions (with opposite characters played forward/backward). Participants watched two runs, each containing 3 clips with 20 seconds of fixation before and after each clip. **(B) Example activation across a run within a language region. Center**: One language ROI (Left AntTemp, green). ss-fROIs were created per subject within language parcels, theory of mind parcels, and conversation spherical parcels. **Within box, left**: Example timecourse for one run of SS-IntDialog, for one participant (light blue), the average of the other participants who heard the exact same version of the run (darker blue), and the average of the participants who heard the opposite version of the run (purple). Background shading indicates when speech was forward (blue) or backward (purple) from the perspective of the held-out participant (opposite for the “other” group: purple is forward and blue is backward).

We had two versions of each clip, such that one group of participants heard one character forward (e.g., Elmo forward and Abby backward) and the other group of participants heard the other character forward (e.g., Abby forward and Elmo backward). The visual information, and the context and social structure of the clip, was preserved (e.g., Elmo is asking Abby about her magic wand). This design allowed us to calculate ISCs between a held-out subject’s timecourse and (1) the average timecourse for other participants who heard the same version of the videos, and (2) the average timecourse for the participants who heard the opposite version of the videos, within ss-fROIs (**Figure 5B**). Comprehensible utterances varied in length from .46 to 34.68 seconds, with a mean(SD) of 3.74(3.84) seconds (**Figure 5B**).

### fMRI Task

Participants watched 1-3-minute dialogue clips of *Sesame Street* in which one character’s audio stream was played forward and the other was played backward. Additional sounds in the video (e.g., blowing bubbles, a crash from something falling) were played forwards. Participants watched the videos and pressed a button on an in-scanner button box when they saw a still image of Elmo appear on the screen immediately after each block. Participants completed 2 runs, each approximately 8 min 52 sec long. Each run contained unique clips, and participants never saw a version of the same clip with the forward/backward streams reversed.

Each run contained 3 clips presented in the same order. Each video was followed by 1.5 seconds of still image attention check (Elmo), 0.5 seconds of a blank screen, then a 22-second fixation block (one run had less total video time, so there was additional rest at the end to reach the 8 min 52 sec acquisition time). There was also a 22-second fixation block at the beginning of the run. Versions of each clip with the opposite character Forward and Backward were counterbalanced between participants (randomly assigned Set A or Set B). 11 participants saw version A, and 9 participants saw version B (1 run from group A was excluded due to participant falling asleep, and one run from group B was excluded due to motion). Run order was randomized for each participant (random sequence 1-2). Transcripts and stimuli features can be found here: https://osf.io/whsb7/

### Univariate Analysis

For first-level modeling, event regressors were created for Forward and Backward speech segments and for the button press response period (when a still image of Elmo appeared on the screen and participants were asked to respond via button press). Each event regressor was defined as a boxcar convolved with a standard double-gamma HRF, with the boxcar defined over the onset to the offset of forward and backward speech segments within the video clips. Statistical analyses were conducted in R, using the average activation per condition within ss-fROIs as described in **General Methods**. Conditions were compared using linear mixed effects models; t-tests used Satterthwaite’s method. We first tested for network-level fixed effects, with ROI and participants modeled as random effects, using: lmer(mean_topvoxels_extracted∼f_or_b+(1|ROI)+(1|participantID), REML = FALSE), where f_or_b was forwards or backwards (coded 1, −1, respectively), and ROI was region of interest within the network. Significance was determined at a level of p<.05 Bonferroni corrected for the two networks tested (left and right language regions). We also examined effects in individual regions: lmer(mean_topvoxels_extracted∼f_or_b+(1|participantID), REML = FALSE).

Significance was determined at a level of p<.05 Bonferroni corrected for the number of ROIs (6 for canonical language regions and 6 for right hemisphere language regions). In exploratory analyses, we also modeled left and right language regions together and tested for interactions with hemisphere, both at a bilateral language network level and in individual regions, coding for left or right (coded 1,-1).

### Intersubject Correlation Analysis

For the SS-IntDialog task, each participant saw two runs, each of which contained three different video clips (in the same order within a run). Half the participants saw version A, and half of the participants saw version B of these runs (same videos, different audio streams). That is, if Elmo was speaking forward in the first clip in Run 1 version A, Elmo spoke backward in the first clip in Run 1 version B. We performed ISC analyses across the entire run, including the rest blocks between clips. ISC analyses were performed using in-lab scripts modeled after the tutorials in https://naturalistic-data.org/ (Chang et al., 2020). The preprocessed data were smoothed with a 6mm kernel, and then denoised using a GLM (6 realignment parameters, their squares, their derivatives, and squared derivatives), with outliers excluded using a dummy code, and average CSF activity and linear and quadratic trends regressed out. The timecourse was z-transformed to be centered at 0.

First, we extracted the timecourse per participant, per run for each language ss-fROI (defined as specified in **General Methods**, using the auditory language localizer). Using a leave-one-subject out approach, we calculated the correlation between the held-out subject’s timecourse (i.e., the average response of that subject across all 100 voxels in that ROI) and (1) the average timecourse of the remaining participants who watched the same version of the stimuli, and (2) the average timecourse of the participants who watched the opposite version of the stimuli, for each language region. Next, we did the same analyses using the extracted timecourses per participant, per run for each of the ToM ss-fROIs. Finally, we repeated the same analysis with the extracted timecourses per participant, per run for each conversation ss-fROI, defined as the top 100 voxels for the [Forward Dialogue>Forward Monologue]>[Backward Dialogue>Backward Monologue] interaction contrast within 10mm spheres centered at the center of gravity point for each significant cluster in the group map (**Supplementary Table 2**).

Statistical analyses were conducted in R. Within each region, one-sample two-tailed t-tests were conducted to determine whether Within-Group and Between-Group correlations differed from 0. Paired t-tests were used to determine if Within-Group correlations differed from Between-Group correlations within each region. Significance was determined at a level of p<.05 Bonferroni corrected the number of ROIs per network (6 for left-hemisphere language, 6 for right-hemisphere language, and 7 for ToM). To test whether Within-Group correlations were higher than Between-Group correlations within each network, we used linear mixed effects models with ROI and participants modeled as random effects: lmer(z_correlation∼w_or_b +(1|participantID)+(1|ROI), REML = FALSE), where w_or_b was within-group or between-group (coded 1, −1), and ROI was region of interest within the network. To test for an interaction with hemisphere, we included both left and right language ROIs within the same model: lmer(z_correlation∼w_or_b*l_or_r_roi +(1|participantID)+(1|ROI), REML = FALSE), where l_or_r_roi was left or right (coded 1, −1). We also checked for interactions with hemisphere in individual ROIs: lmer(z_correlation∼w_or_b*l_or_r +(1|participantID, REML = FALSE).

## Results

### Univariate response to forward and backward speech

By modeling the onset and offset of each utterance within the extended SS-IntDialog dialogues, we replicated the robust response to forward utterances, and the very low response to backward utterances, in the canonical left-hemisphere language network (Forward>Backward: Est.=1.27, S.E.=0.08, t-value=15.53, corrected p-value<.001), as well as in individual left-hemisphere language regions (**Figure 6**, **Supplementary Table 3**). Right hemisphere homologues of language regions likewise responded more to forward than backward speech at a network level (Forward>Backward: Est.=0.70, S.E.=0.09, t-value=7.46, corrected p-value<.001), and at the level of individual regions with the exception of right AngG (**Supplementary** Figure 2, **Supplementary Table 2**). When both right and left hemisphere language regions were included in the model, there was a main effect of comprehensibility (Forward>Backward: Est.=0.99, S.E.=0.06, t-value=15.18, corrected p-value<.001) and an interaction between comprehensibility and hemisphere (Forward>Backward*Left>Right: Est.=0.29, S.E.=0.06, t-value=4.39, uncorrected p-value<.001). Thus, canonical left-hemisphere language regions (and right-hemisphere homologues) responded robustly to the timing of comprehensible utterances within the audio stream.

**Figure 6:**
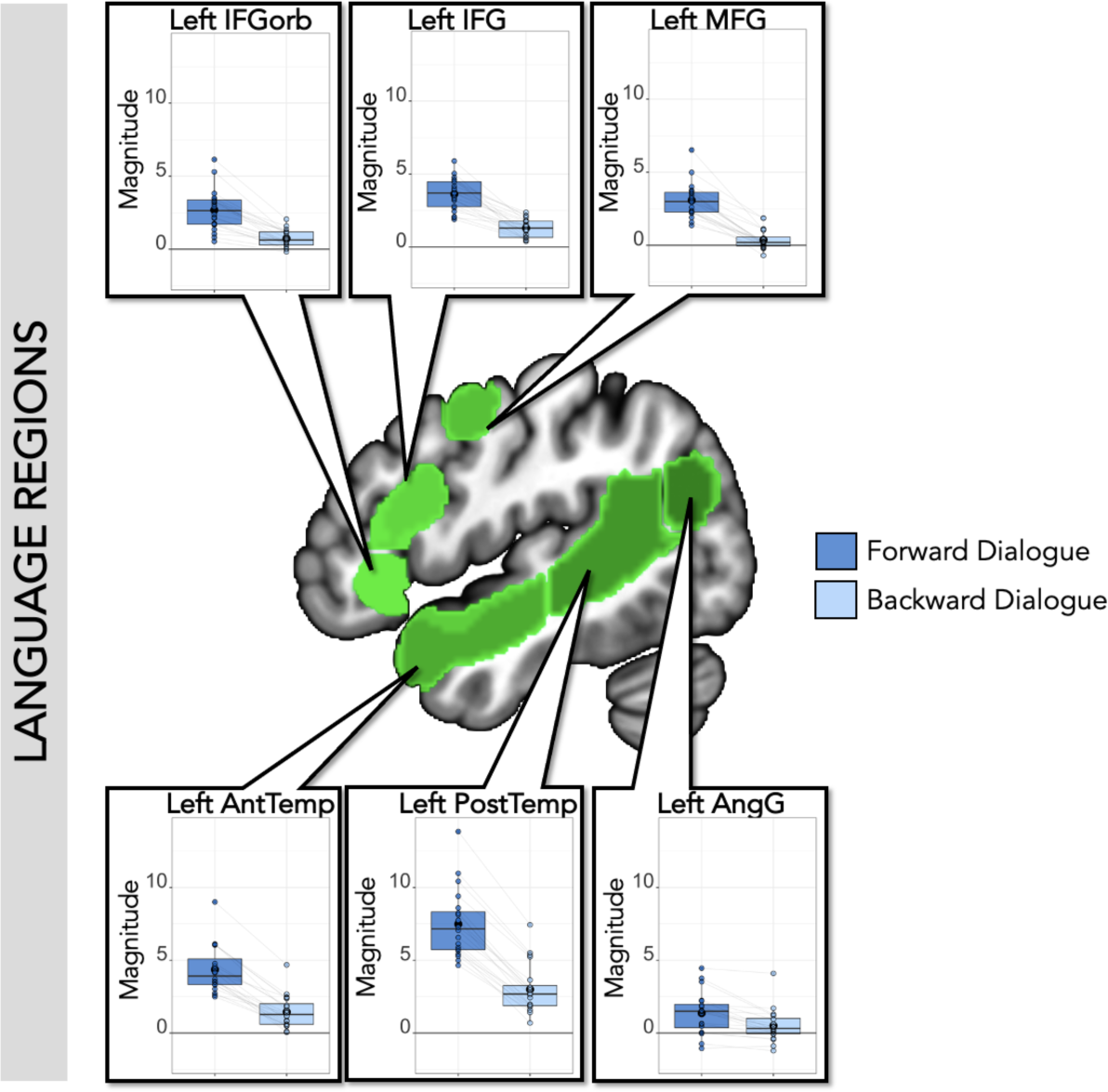
SS-IntDialog average magnitude by condition within language regions. **Center**: Left hemisphere language parcels overlaid on template brain (green; parcels include left IFGorb, IFG, MFG, AntTemp, PostTemp, and AngG from https://evlab.mit.edu/funcloc/). **Panels**: Average response magnitude per individual for each condition in the SS-IntDialog task was extracted from subject-specific functional regions of interest for language (blue: Forward Dialogue; light blue: Backward Dialogue). All regions responded more to Forward than Backward speech. Each individual’s datapoints are connected by light gray lines.

### Timecourse of response to dialogue videos in language regions

The key analysis of Experiment Task 2 used intersubject correlations (ISCs) to test identify the stimulus-driven structure of the regions’ timecourses. The timecourse of response in canonical left hemisphere language regions was correlated across participants who saw the same version of the extended dialogue, with the same character’s speech played forward (Within-Group Correlations: M(SD)=0.41(0.17), one-sample t-test against 0 was significant (95% confidence interval: 0.38-0.44, t-value= 26.79, p-value<.001); for all regions, one-sample t-test against 0 was significant; **Figure 7**; **Table 4**). Thus, the short comprehensible utterances within these dialogues drove reliable responses, consistently across participants. In contrast, when comparing the timecourse to participants hearing the opposite character’s speech played forward, there was little to no correlation in canonical left-hemisphere language regions (average Between-Group Correlations: M(SD)=0.04(0.08), one-sample t-test against 0 was significant (95% confidence interval: 0.02-0.05, t-value=4.68, p-value<.001); one-sample t-test testing for greater than 0 was not significant in individual regions except left PostTemp and AngG; **Figure 7**; **Table 5**). Even the significant Between-Group correlations in PostTemp and AngG were weak (PostTemp r=0.07, AngG r=0.08) and were below zero for some participants. In the network, and in every individual region, ISCs were much higher within than between groups (network, Within>Between: Est.=0.18, S.E.=0.007, t-value=26.13, corrected p-value<.001; for individual regions **Table 5**). These results suggests that reliable temporal structure in these regions was driven by language comprehensibility, and not by the visual and abstract semantic structure of the dialogues preserved between the groups (e.g., the sequence of visual images, the topic of the conversation, etc.).

**Figure 7:**
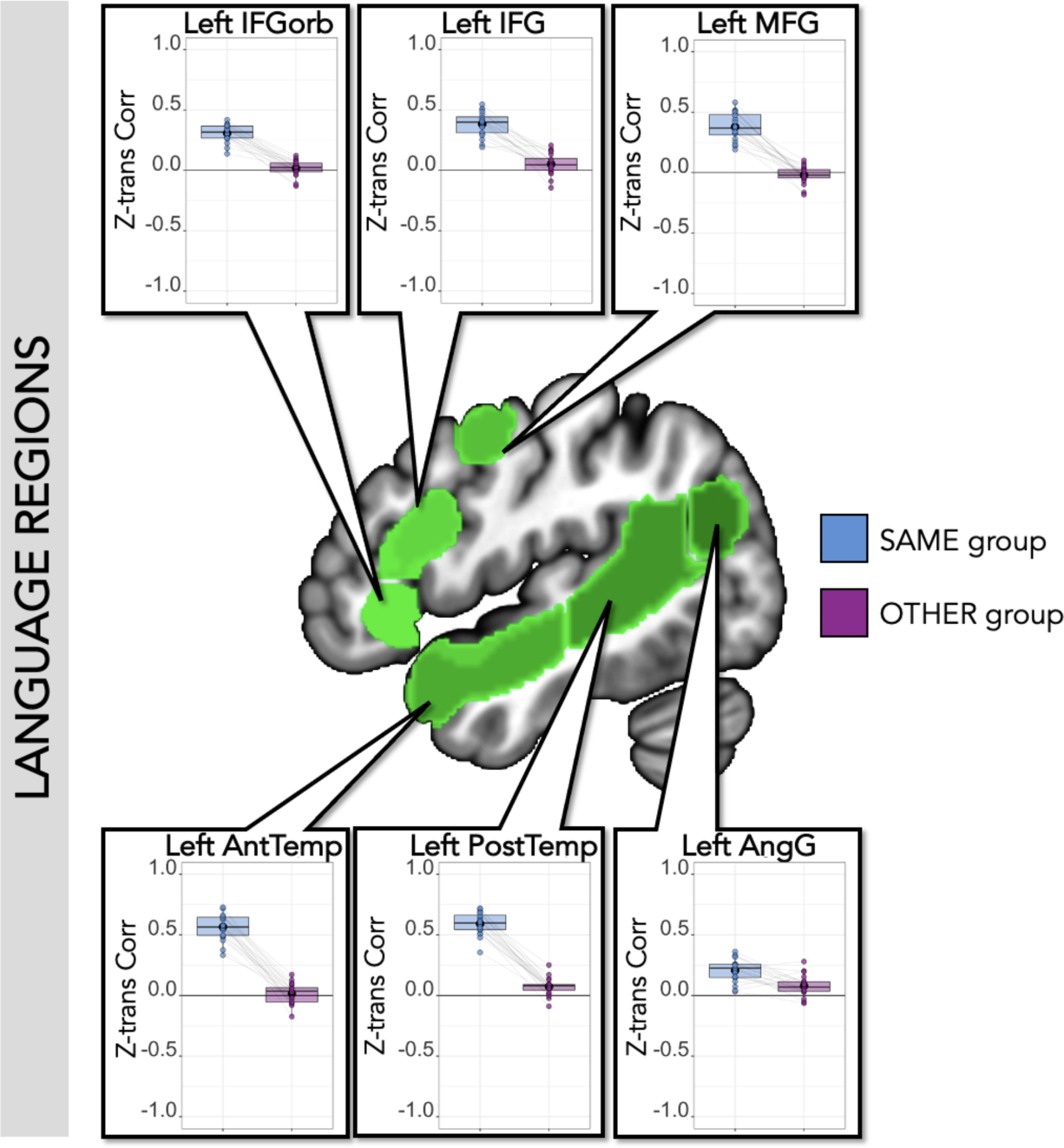
SS-IntDialog correlations within language regions. **Center**: Left hemisphere language parcels overlaid on template brain (green; parcels include left IFGorb, IFG, MFG, AntTemp, PostTemp, and AngG from https://evlab.mit.edu/funcloc/). **Panels**: Average z-transformed Pearson’s correlation between each held-out subject’s timecourse within each ss-fROI and the average timecourse of the remaining participants who viewed and listened to the same version of the stimuli (blue) and the average of the participants who heard the opposite audio stream (purple). Each individual’s datapoints are connected by light gray lines. Within-group correlations were higher than between-group correlations in all regions.

**Table 4:** SS-IntDialog timecourse correlations within language regions. Average z-transformed Pearson’s correlations between each held-out subject and the average of the rest of the group that heard the same version of the clips (within-group) and the average of the group that heard the opposite version of the clips. One-sample t-test shows significance test for two-tailed t-test against 0 (uncorrected p-values reported). Paired t-test shows that there were higher within-group than between-group correlations for each canonical language region (uncorrected p-values reported). * indicates significance level p<.05, Bonferroni corrected for 6 ROIs (p<.0083)

| Language Regions |  |  |  |
| --- | --- | --- | --- |
| ROI | Within-Group Correlation | Between-Group Correlation | Paired T-test (W v. B) |
| Left IFGorb | M(SD) = 0.31(0.07);<br>range = 0.14-0.42<br><u>One-sample t-test:</u><br>t-value = 19.02, $p < .001$ * | M(SD) = 0.02(0.06);<br>range = -0.13-0.12<br><u>One-sample t-test:</u><br>t-value = 1.19, p-value = 0.25 | $t = 14.32$, $p < .001$ * |
| Left IFG | M(SD) = 0.38(0.10);<br>range = 0.19-0.55<br><u>One-sample t-test:</u><br>t-value = 17.30, $p < .001$ * | M(SD) = .05(0.09);<br>range = -.14-.21<br><u>One-sample t-test:</u><br>t-value = 2.59, p-value = 0.02 | $t = 10.02$, $p < .001$ * |
| Left MFG | M(SD) = 0.38(0.11);<br>range = 0.19-0.58<br><u>One-sample t-test:</u><br>t-value = 15.07, $p < .001$ * | M(SD) = -0.02(0.07);<br>range = -0.18-0.10<br><u>One-sample t-test:</u><br>t-value = -1.27, p-value = 0.22 | $t = 11.95$, $p < .001$ * |
| Left AntTemp | M(SD) = 0.57(0.11);<br>range = 0.33-0.73<br><u>One-sample t-test:</u><br>t-value = 24.01, $p < .001$ * | M(SD) = 0.02(0.08);<br>range = -0.17-0.17<br><u>One-sample t-test:</u><br>t-value = 0.81, p-value = 0.43 | $t = 18.32$, $p < .001$ * |
| Left PostTemp | M(SD) = 0.59(0.09);<br>range = 0.36-0.72<br><u>One-sample t-test:</u><br>t-value = 28.96, $p < .001$ * | M(SD) = 0.07(0.07);<br>range = -0.09-0.25<br><u>One-sample t-test:</u><br>t-value = 4.53, $p < .001$ * | $t = 22.36$, $p < .001$ * |
| Left AngG | M(SD) = 0.21(0.09);<br>range = 0.03-0.36<br><u>One-sample t-test:</u><br>t-value = 9.77, $p < .001$ * | M(SD) = 0.08(0.08);<br>range = -0.06-0.28<br><u>One-sample t-test:</u><br>t-value = 4.20, $p < .001$ * | $t = 4.73$, $p < .001$ * |

**Table 5:**
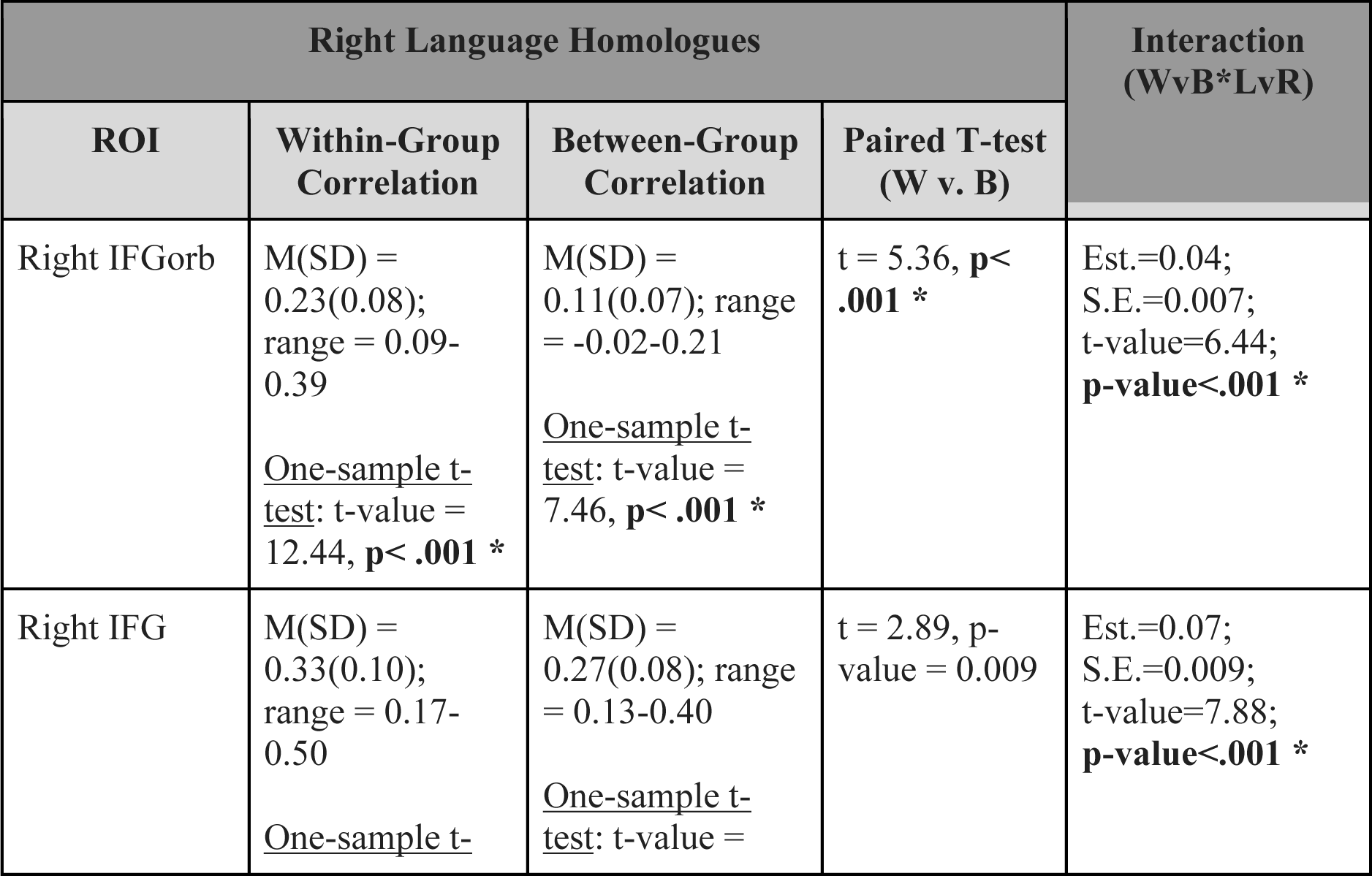
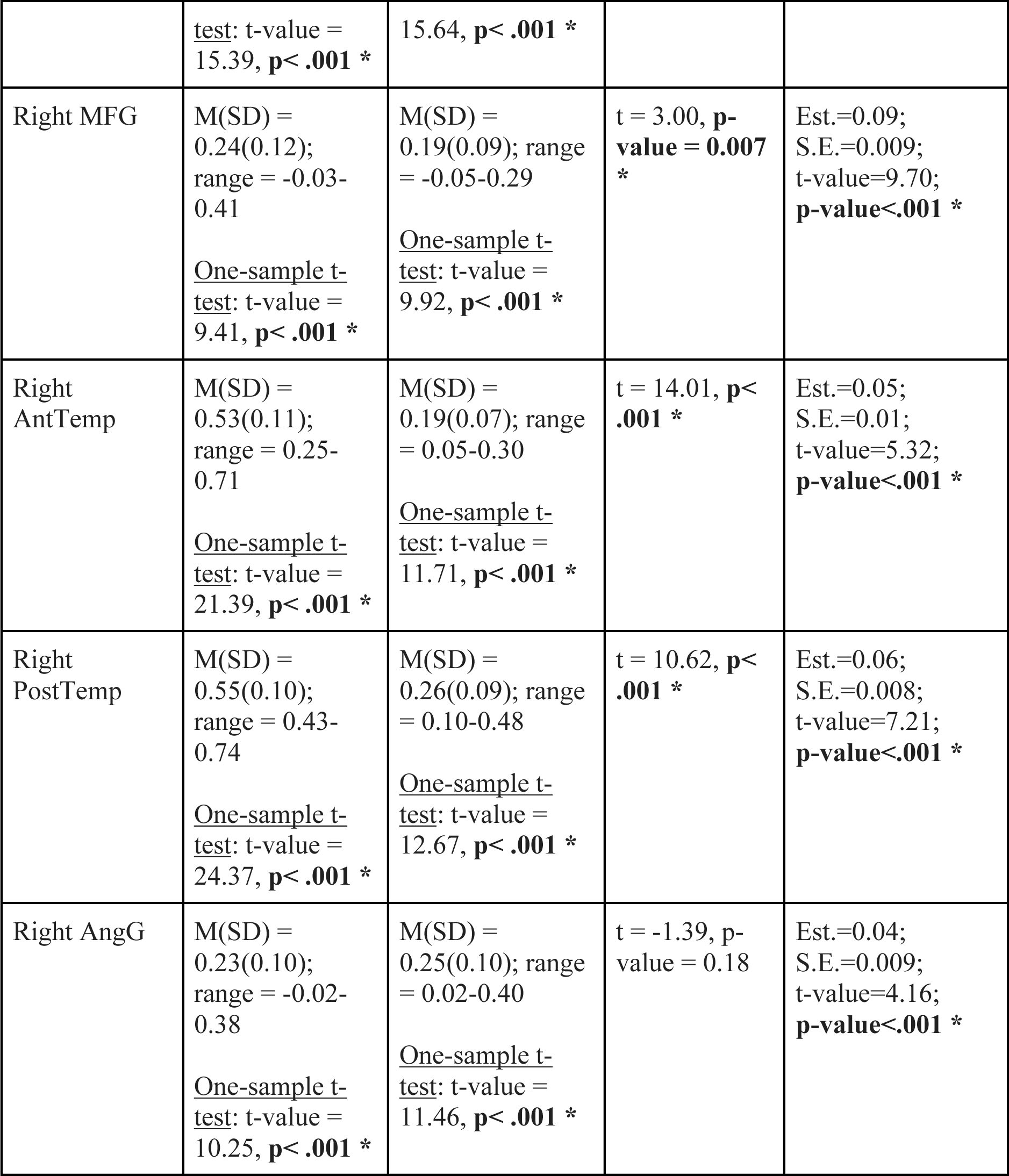
SS-IntDialog timecourse correlations within right language region homologues. Average z-transformed Pearson’s correlations between each held-out subject and the average of the rest of the group that heard the same version of the clips (within-group) and the average of the group that heard the opposite version of the clips. One-sample t-test shows significance test for two-tailed t-test against 0 (uncorrected p-values reported). Paired t-test shows that there were higher within-group than between-group correlations for each right hemisphere language region except right IFG and AngG (uncorrected p-values reported). Interaction (WvB*LvR) column shows results for the interaction between within vs between and left vs right hemisphere from the model: lmer(z_correlation∼w_or_b*l_or_r_roi+(1|participantID), REML = FALSE) * indicates significance level p<.05, Bonferroni corrected for 6 ROIs (p<.0083)

### Timecourse of response to dialogue videos in other regions

As a point of comparison to left-hemisphere cortical language regions, we also examined timecourse similarity between groups in right language homologues, ToM regions, and “comprehensible dialogue” regions identified by the exploratory whole-brain analysis described in **Experimental Task 1**. In many cases, unlike in left-hemisphere language regions, the dialogue videos *did* drive shared temporal structure not only within, but also across participant groups. In right hemisphere language homologues, both the within-group and between-group correlations were positive for all regions (Within-Group: M(SD)=0.35(0.17), 95% confidence interval: 0.32-0.38, t-test against 0: t-value=22.67, p-value<.001; Between-Group: M(SD)=0.21(0.10), 95% confidence interval: 0.19-0.23, t-test against 0: t-value=23.49, p-value<.001; **Figure 8A**; **Table 5**), though the within-group correlations were higher than the between-group correlations across the network (Within>Between: Est.=0.07, S.E.=0.007, t-value=10.43, corrected p-value<.001) and for all regions except right IFG and AngG. When both right and left hemisphere language regions were included in the same model, there was a main effect of group comparison (Within>Between: Est.=0.13, S.E.=0.005, t-value=26.08; p-value<.001), and critically, there was an interaction between group comparison and hemisphere (Within>Between*Left>Right: Est.=0.06, S.E.=0.005, t-value=11.75; corrected p-value<.001). This interaction was also significant in each region (**Table 5**), showing that the difference between the within-group and between-group correlations was greater in each left hemisphere language region than its right hemisphere homologue. Overall, between-group correlations were higher for right-hemisphere than left-hemisphere regions (Left>Right: Est.=-.09, S.E.=.01, t-value=-6.39, p-value<.001).

**Figure 8:**
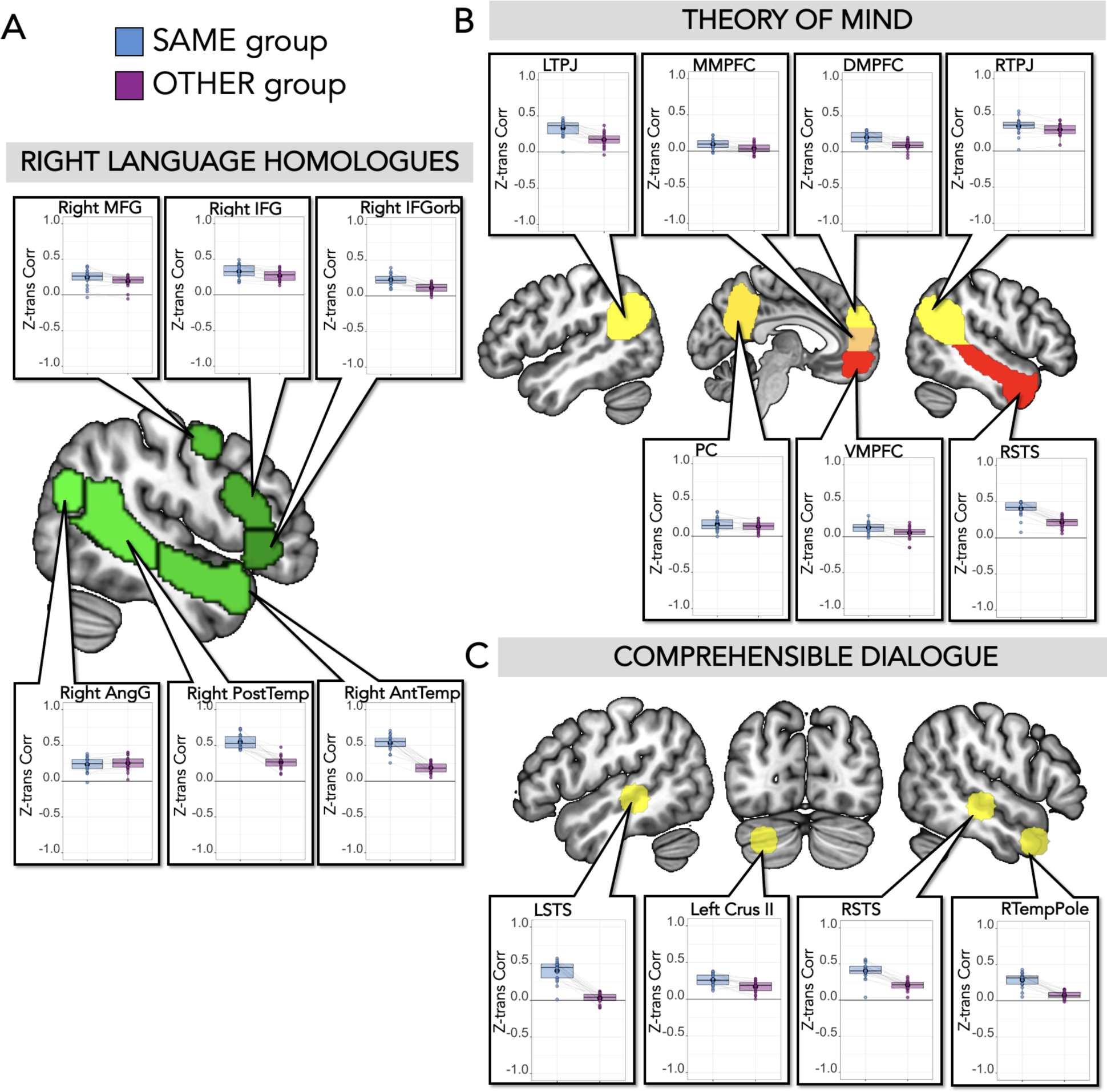
SS-IntDialog correlations within right language homologues, theory of mind regions, and comprehensible dialogue regions. **(A) Right Language Homologues. Center:** Right hemisphere language parcels overlaid on template brain (green; parcels include right IFGorb, IFG, MFG, AntTemp, PostTemp, and AngG from https://evlab.mit.edu/funcloc/). **Panels**: Average z-transformed Pearson’s correlation between each held-out subject’s timecourse within each ss-fROI and the average timecourse of the remaining participants who viewed and listened to the same version of the stimuli (blue) and the average of the participants who heard the opposite audio stream (purple), averaged across two runs. Each individual’s datapoints are connected by light gray lines. **(B) Theory of Mind. Center**: Theory of mind parcels overlaid on template brain (parcels include LTPJ, MMPFC, DMPFC, RTPJ, PC, VMPFC, and RSTS from (Dufour et al., 2013)). **Panels**: For all regions except MMPFC and PC, the within-group correlations were higher than the between-group correlations. Each individual’s datapoints are connected by light gray lines. **(C) Comprehensible Dialogue ROIs. Top**: 10mm spherical search spaces (yellow) based on COG coordinates from clusters identified in the group whole-brain analysis for comprehensible dialogue ([Forward Dialogue>Forward Monologue]>[Backward Dialogue>Backward Monologue]). **Panels**: For all regions, the within-group correlations were higher than the between-group correlations. Each individual’s datapoints are connected by light gray lines.

Brain regions defined by the independent ToM localizer also showed significant positive correlations in the timecourse of responses both within-group (Within-Group: M(SD)=0.24(0.15), 95% confidence interval: 0.21-0.26, t-test against 0: t-value=19.26, p-value<.001; for all regions, one-sample t-test against 0 was significant) and between-groups (Between-Group: M(SD)=0.14(0.12), 95% confidence interval: 0.12-0.16, t-test against 0: t-value=14.36, p-value<.001; for all regions except VMPFC and MMPFC, one-sample t-test against 0 was significant). There were again higher within-group correlations than between-group correlations across ToM network (Within>Between: Est.=0.05, S.E.=0.005, t-value=10.36, corrected p-value<.001), and for all regions except MMPFC and PC (**Figure 8B**; **Table 6**).

**Table 6:**
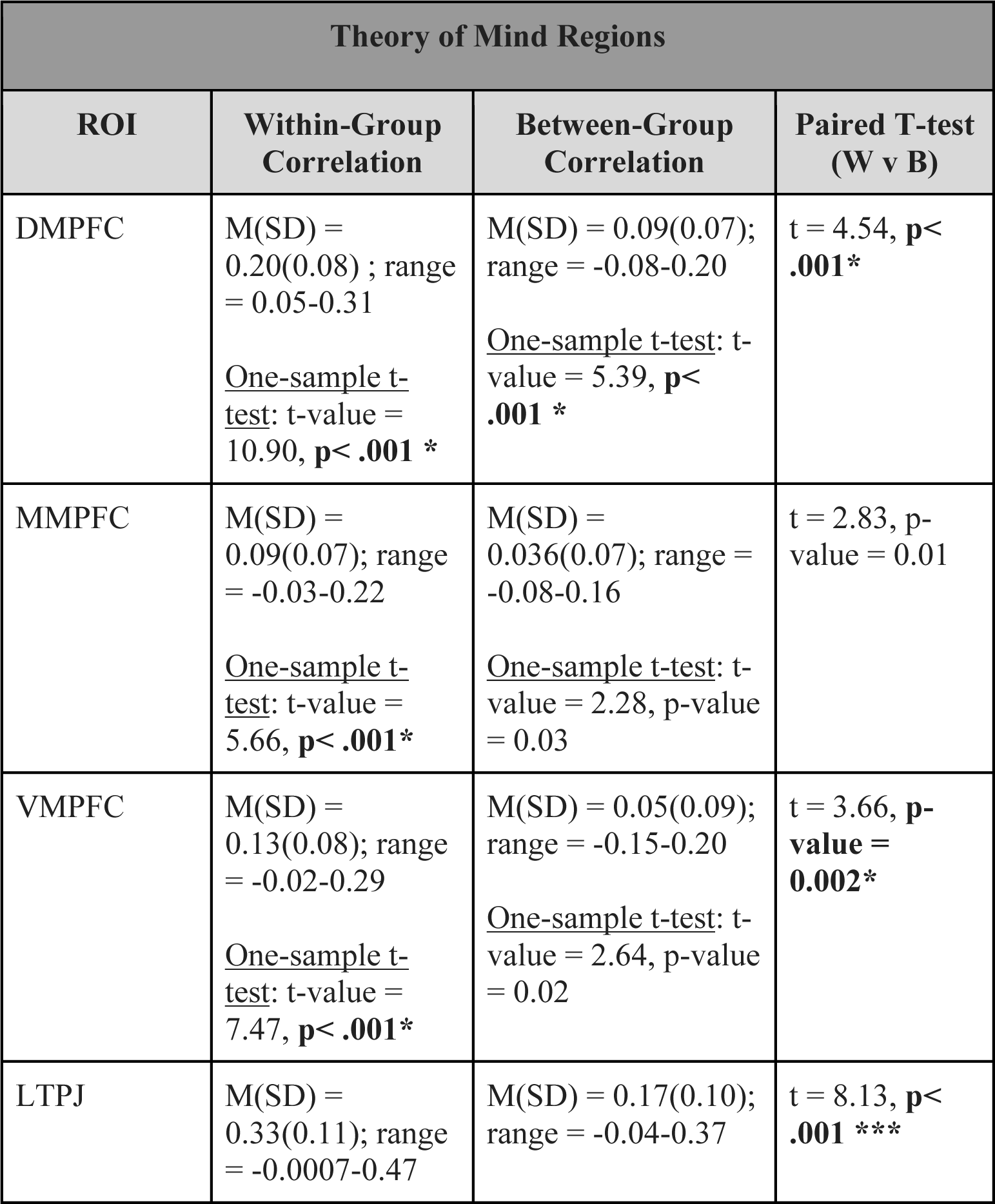
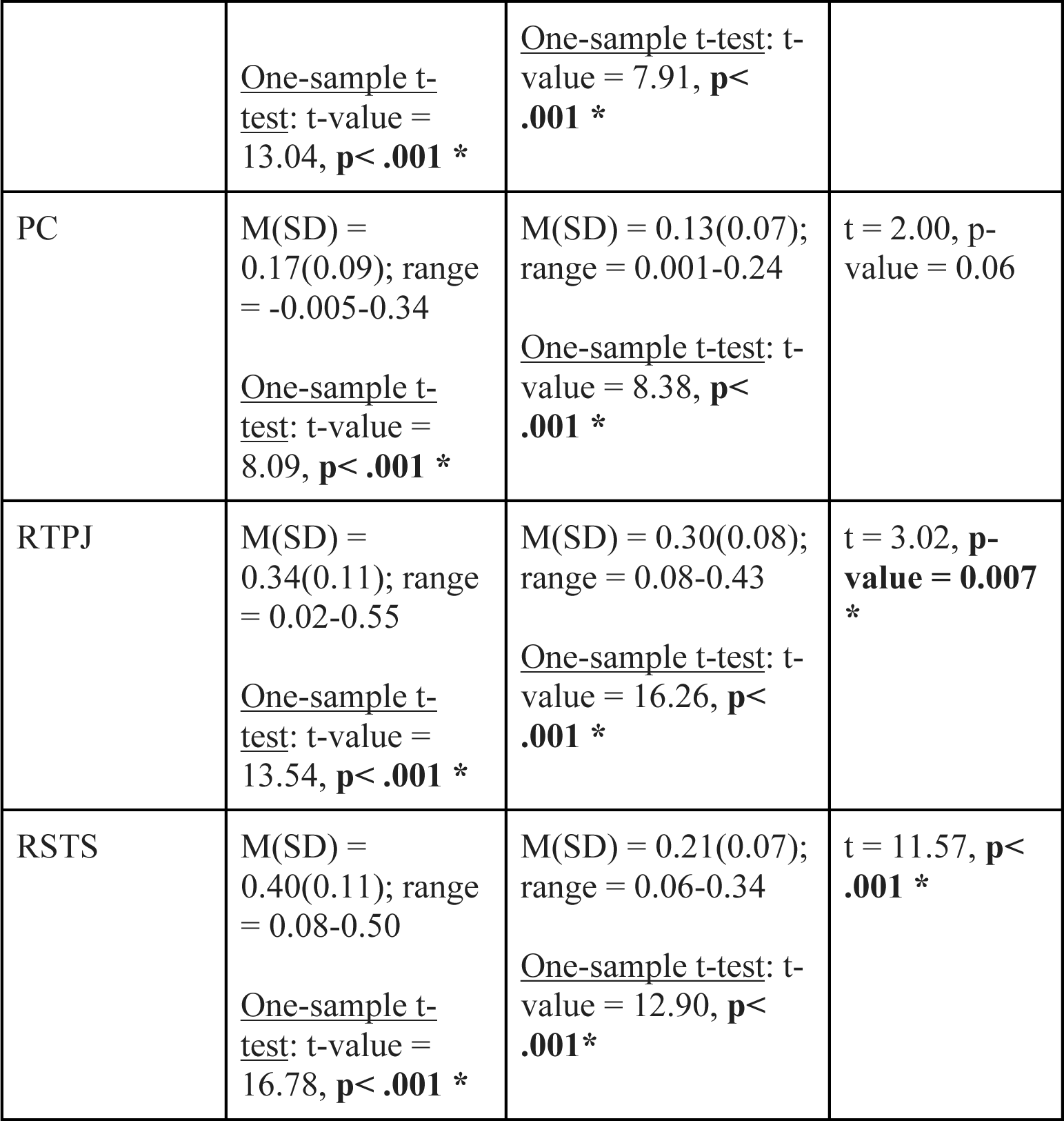
SS-IntDialog timecourse correlations within theory of mind regions. Average z-transformed Pearson’s correlations between each held-out subject and the average of the rest of the group that heard the same version of the clips (within-group) and the average of the group that heard the opposite version of the clips. One-sample t-test shows significance test for two-tailed t-test against 0 (uncorrected p-values). Paired t-test shows that there were higher within-group than between-group correlations for each ToM region except MMPFC and PC (uncorrected p-values). * indicates significance level p<.05, Bonferroni corrected for 7 ROIs (p<.0071)

When both left-hemisphere language regions and ToM regions were included in the same model, there was a main effect of group comparison (Within>Between: Est.=.12, S.E.= 0.004, t-value=28.29; p-value<.001), and critically, there was an interaction between group comparison and network (Within>Between*Left_Language>ToM: Est.= 0.07, S.E.= 0.004, t-value=16.62; p-value<.001). Between-group correlations were higher in ToM regions than left-hemisphere language regions (Left_Language>ToM: Est.=-0.05, S.E.=0.02, t-value=-2.78, p-value=.02).

Finally, the brain regions in right STS, right temporal pole, left STS, and left Crus II (cerebellum), identified as specifically responsive to comprehensible dialogue in Experimental Task 1, similarly showed correlated timecourses both within-groups and between-groups (with the exception of left STS for the between-groups correlation; **Figure 8C**; **Table 7**). Thus, the preserved visual and abstract semantic structure of the dialogue drove reliable cortical responses outside of the canonical left hemisphere language regions.

**Table 7:** SS-IntDialog timecourse correlations within conversation regions. Average z-transformed correlations between each held-out subject and the average of the rest of the group that heard the same version of the clips (within-group) and the average of the group that heard the opposite version of the clips. One-sample t-test shows significance test for two-tailed t-test against 0 (uncorrected). Paired t-test shows higher within-group than between-group correlations within each region. No p-values are reported since analyses were exploratory.

| Comprehensible Dialogue Regions |  |  |  |
| --- | --- | --- | --- |
| ROI | Within-Group Correlation | Between-Group Correlation | Paired T-test (W v B) |
| RTempPole | M(SD) = 0.29(0.10); range = 0.05-0.43<br><u>One-sample t-test</u>: t-value = 13.34 | M(SD) = 0.08(0.05); range = -0.01-0.16<br><u>One-sample t-test</u>: t-value = 6.79 | t = 10.28 |
| RSTS | M(SD) = 0.40(0.12); range = 0.04-0.56<br><u>One-sample t-test</u>: t-value = 15.45 | M(SD) = 0.21(0.06); range = 0.04-0.32<br><u>One-sample t-test</u>: t-value = 14.21 | t = 10.01 |
| LCere | M(SD) = 0.26(0.08); range = 0.12-0.38<br><u>One-sample t-test</u>: t-value = 14.56 | M(SD) = 0.17(0.08); range = 0.003-0.28<br><u>One-sample t-test</u>: t-value = 9.69 | t = 4.0738 |
| LSTS | M(SD) = 0.40(0.14); range = 0.01-0.57<br><u>One-sample t-test</u>: t-value = 12.95 | M(SD) = 0.03(0.07); range = -0.10-0.12<br><u>One-sample t-test</u>: t-value = 1.87 | t = 10.16 |

### Summary

Canonical left-hemisphere cortical language regions responded robustly to the local structure of comprehensible utterances within a continuous auditory stream. Furthermore, the timecourses of activity in canonical left-hemisphere language regions were similar among individuals listening to the same language input, but not among individuals watching the same videos while listening to the reciprocal language input (i.e., the opposite speaker in the dialogue). The strikingly low correlations in the response of canonical left-hemisphere language regions when listening to opposite auditory streams suggest that language regions are insensitive to the larger social, contextual and visual structure of the dialogue videos. On the other hand, theory of mind regions and right hemisphere homologues of language regions were correlated even when participants were listening to the opposite auditory streams.

## General Discussion

Multimodal dialogue is a useful boundary test case for probing the scope of processing carried out by canonical left-hemisphere cortical language regions: tracking multiple speakers and perspectives is part of comprehending language in dialogue, but it is not a function that has been attributed to these regions. In two tasks, we manipulated the audio stream of *Sesame Street* videos to create matched segments of naturalistic videos with forward (comprehensible) and backward (incomprehensible) speech. These naturalistic multimodal stimuli thus evoke cognitive processes typically involved in understanding dialogues: alternating speech from two identifiable and distinct speakers interacting in context. We defined three measures of a cortical region’s (in)sensitivity to the dialogue context of linguistic input. First, a region that processes language independent of a dialogue context should respond equally robustly to forward speech whether presented as a monologue or dialogue. Second, it should respond selectively to the comprehensible speech segments in a dialogue that alternates between forward and backwards speech, even within the frequent alternations of dialogue that render some utterances quite short. Third, the reliable (between-participants) stimulus-driven timecourse of response to these alternating dialogue stimuli should be driven only by the timing of the comprehensible speech segments, and not by any other features of the dialogue. By all three of these measures, we find that left hemisphere canonical language regions are insensitive to whether language is in the form of dialogue during passive observation.

### Insensitivity to non-linguistic features of observed dialogue in canonical language regions

We chose two different analytic approaches to test whether language regions are sensitive to any cognitive processes – other than language comprehension – evoked by observing dialogue. Using a block design (Experimental Task 1), we found no differences in the magnitude of neural response to multimodal dialogue versus monologue in the canonical left-hemisphere language network, regardless of whether the speech was comprehensible. These results are consistent with prior evidence that language regions are not sensitive to factors that are language-adjacent, such as reading computer code (Ivanova et al., 2020; Liu et al., 2020). Nor, it seems, do language regions process differences between monologue and dialogue, at least not to a level that was detectable using this fairly blunt approach.

The structure of Experimental Task 2 afforded a complementary perspective to examine dialogue comprehension in language regions. With the longer audiovisual stimuli in Experimental Task 2, we could use ISC analyses (Hasson et al., 2004) to ask: does any feature other than language comprehensibility at the level of individual utterances drive the neural responses in language regions during a dialogue? This approach was again blunt, in that we did not directly manipulate and test specific features of the dialogue, but also sensitive, in that we could simultaneously examine the effects of many sparser features of multimodal dialogue clips. Participants who heard the same audio with each video clip had similar responses to the stimuli in left-hemisphere cortical language regions (i.e., positive within-group correlations within language regions).

Critically, though, participants who watched the same video clips with the reciprocal audio stream showed close to zero correlation in canonical language network activity. Thus, other features of the visually displayed dialogue, other than the comprehensibility of the speech stream, were not reliably tracked by the left-hemisphere language network across participants. Our two analytic approaches, and two task designs, both produced complementary results that suggest canonical left-hemisphere cortical language regions are not sensitive to features of dialogue other than the occurrence of comprehensible utterances. This insensitivity is consistent with other evidence that language regions are sensitive to relatively local linguistic features, and with evidence that canonical language regions have fairly short temporal receptive windows (Blank & Fedorenko, 2020; Hasson et al., 2008).

### Sensitivity to non-linguistic features of observed dialogue outside language regions

By contrast to canonical left-hemisphere cortical language regions, there was evidence of sensitivity to dialogue in the other individually functionally-defined regions: ToM regions and right hemisphere homologues of language regions. Like the left-hemisphere language network, the ToM network responded more to forward than backward speech in Experimental Task 1 – but unlike the left-hemisphere language network, the ToM network also responded more to dialogue than monologue, and had an interaction between Forward>Backward and Dialogue>Monologue. This pattern appeared in some individual ToM regions (see **Figure 3B**). Note, however, that the three-way interaction between experimental conditions and network (ToM versus Language) was not significant in Experimental Task 1. In Experimental Task 2, the ToM network and most individual ToM regions showed positive correlations in the timecourses of between-group participants, who saw the same videos with reciprocal audio streams. In this case, the difference between networks did produce a significant interaction.

Unlike language, which was experimentally manipulated by the forward and backward conditions, we did not directly manipulate or vary social demands in the clips. Thus, while we know that we were examining responses in ToM regions because we localized them using a separate, well-validated task, we can only speculate on why these regions respond most in the dialogue condition. In experiments using single source texts, ToM regions respond selectively to stimuli that describe or imply contrasting beliefs, knowledge or emotions, between characters or over time (Dodell-Feder et al., 2011; Saxe & Kanwisher, 2003; Saxe & Powell, 2006). However, there was no difference in the occurrence of terms explicitly referring to mental states between the dialogue and monologue stimuli in our task. Instead, the ToM demands likely arose from the social interaction in the dialogues. Naturalistic dialogue often implies differences of perspective, both through information carried by language as well as non-linguistic audio and visual cues, as speakers use utterances to show how a prior utterance was or was not understood. For example, dialogue videos showed Grover trying to sell Kermit a comb, Abby teaching Elmo how to use a magic wand, Elmo teaching Abby how to drink from a straw, Ernie watching Bert wrap a present, and Grover teaching Elmo how to deliver a telegram. In each of these cases, observers may spontaneously consider the differences in the two characters’ knowledge and desires, and how those differences change over time, while viewing the videos. Even in Experimental Task 2, when only one character’s speech was comprehensible, the overall context of the video conveyed the social interaction and the difference of perspectives. For example, whether a participant hears Grover’s part of the dialogue (“Oh Sir, I’m so sorry, you are absolutely correct. The frame is a circle and your painting is a square.”) or hears Mr. Johnson’s part (“And now you brought … a TRIANGLE!”), the difference in perspectives between the characters and the change in their mental states over time are apparent. The social and narrative structure of the dialogue, over slower timescales than the individual utterances, thus likely explain the between-group correlations in the ISCs in ToM regions (Hasson et al., 2008; Redcay & Moraczewski, 2020).

We hypothesize that processing observed dialogue in natural social contexts requires integration between complementary language and ToM processing. Language and ToM regions show robust functional differences, yet rich naturalistic contexts like dialogue typically draw on both types of processing. There are many other examples of simultaneous complementary activity in ToM and language networks. For instance, ToM regions and language regions can both respond strongly to stories about false beliefs, but for different reasons – mental state reasoning in ToM regions and linguistic demands in language regions (Shain et al., 2022). Indeed, across a wide variety of stimuli that vary linguistic and ToM demands – including dialogue –language regions track linguistic stimuli, and ToM regions track stimuli with mental state content (Paunov et al., 2022). Yet there is a special relationship between the two networks – language regions and ToM regions have increased synchronized activity during story comprehension (Paunov et al., 2019).

The other set of regions we examined was right language homologues. This is a particularly interesting point of comparison to the left-hemisphere language regions, since we functionally localized these regions in the same way -by using the independent language localizer task, and selecting top voxels for comprehensible>incomprehensible (specifically, Intact>Degraded) language. Unlike left hemisphere regions, right hemisphere language regions (especially right AntTemp and PostTemp) showed an overall stronger response to dialogue versus monologue, and all of the right hemisphere language homologues showed significant correlations between participants listening to opposite versions of the audio stream in Experimental Task 2. The difference between left and right hemisphere regions in responses to dialogue showed a significant hemisphere by function interaction in Experiment 2 (**Figure 8A**). These results are consistent with prior studies of the right lateralization of social processing (Rajimehr et al., 2022). Previous work has shown that right hemisphere damage can make it more difficult for individuals to make inferences from discourse (Beeman, 1993). Other studies have also demonstrated that pragmatic and social aspects of language may be processed by regions in the right hemisphere. For instance, processing emotional prosody has been shown to be right lateralized (Friederici, 2011; Frühholz et al., 2012; Ross & Monnot, 2008; Seydell-Greenwald et al., 2020), and regions responsive to prosody differences are distinct from language regions, even among individuals with large perinatal strokes in the left hemisphere whose language regions are located in the right hemisphere (Newport et al., 2022). Future work could specifically probe the features of dialogue that drive responses in these regions during complex, real-world language processing.

Finally, in addition to looking within specific regions, we also examined responses across the whole brain to determine where comprehensible dialogue specifically led to higher activation. In exploratory analyses, significant clusters were identified in right temporal pole, right STS, left STS, and left Crus II of the cerebellum (though note that none of these survived correction for multiple comparisons). While these results are exploratory, these regions may be useful targets for future studies on dialogue comprehension. Part of right temporal pole, for example, has long been thought to be involved in social and emotional processing, among other higher level cognitive functions (Herlin et al., 2021; Olson et al., 2007; Pehrs et al., 2017; Wakusawa et al., 2007), and parts of the cerebellum are also involved in language and social cognition (D’Mello & Stoodley, 2015; Stoodley, 2012; Van Overwalle et al., 2014), including in Crus II as we observed (see (Van Overwalle et al., 2020) for a recent meta-analysis). Thus, it is plausible that the clusters we identified in these regions might be meaningful subregions involved in dialogue comprehension. Right STS in particular is a key region supporting social interaction processing, though it is important to note that this is a large region with multiple subregions subserving different functions (Deen et al., 2015). Parts of STS respond to visual social interactions (Walbrin et al., 2018; Walbrin & Koldewyn, 2019), and in particular, a specific part of posterior STS responds to interactions between agents (Isik et al., 2017). Using a naturalistic dataset, part of STS was shown to be selective for interaction, separate from ToM (Lee Masson & Isik, 2021). Other evidence points to additional roles of STS regions in social processing, such as directing attention (Materna et al., 2008) and processing prosody (Wildgruber et al., 2006). Given the interactive nature of the dialogue stimuli, it makes sense that part of STS might be involved in processing the social interaction that occurs in a comprehensible dialogue, either as a subregion that responds to both social interaction and voices, or because the content of the language enhances the perception of an interaction.

### Limitations and Future Directions

Both of the current Experimental Tasks used as stimuli experimentally manipulated audiovisual clips from professionally produced episodes of the television show, *Sesame Street*. There are complementary strengths and limitations of this approach.

First, to measure responses to comprehensible speech, we manipulated the audio stream of each video, reversing utterances to render them incomprehensible. Backward speech is a commonly used control condition in studies of language (e.g., (Bedny et al., 2011; Moore-Parks et al., 2010; Olulade et al., 2020)). In the context of the current study, reversing individual utterances within dialogue videos meant that the recognizable voice of each character and the duration of each utterance were preserved. On the other hand, backward speech is not matched to forward speech in mid-level auditory statistics or phonemes. Thus, the contrast of forward versus backward speech varied the auditory structure of the speech as well as the semantic and syntactic content of the language. Future studies could use acoustically degraded speech (Overath et al., 2015; Stoppelman et al., 2013) or dubbed foreign speech (Schlosser et al., 1998), as control conditions in a similar design.

Second, to measure responses to dialogue, we selected pre-existing video clips of dialogues and monologues. The benefit of using pre-existing commercially-produced videos is that they are highly engaging. Because the clips were drawn from *Sesame Street,* both dialogues and monologues used simple syntax and vocabulary. The average age of acquisition of the words in dialogues and monologues were matched, and before age three years. On the other hand, the linguistic complexity of the stimuli did vary between conditions, with longer average utterances in the monologue clips (**Supplementary** Figure 1). Thus, the contrast of dialogue versus monologue varies both the interaction of speakers and the complexity of the language within each speaker. Future studies could use purpose-generated stimuli in which precisely matched (or exactly the same) sentences are spoken either by a single speaker or by multiple speakers.

Third, the current analyses focus on maximal contrasts between dialogue and monologue, and do not address what drives regions’ responses within either condition. In particular, for right hemisphere homologues and ToM regions, the current analyses cannot identify which specific features of the dialogue drive the greater responses (in Experimental Task 1) or the correlated responses (in Experimental Task 2). Dialogues included more distinct types of speech acts (e.g., questions, backchanneling) and more sentence fragments. There may be other linguistic differences that we have not quantified, for example in the ambiguity of pronoun reference. A complementary analysis approach, such as item-analysis (e.g., (Dodell-Feder et al., 2011)) or encoding models (Naselaris et al., 2011) could test more specific hypotheses. Transcriptions of all of the stimuli, and the raw neural responses, are shared along with this manuscript to make such analyses possible.

Fourth, using videos of dialogue allowed us to investigate neural processes associated with observing dialogue, but not those required to participate in ongoing dialogue. Most theories of dialogue processing concern the special cognitive demands on dialogue participants (e.g., (Bögels & Levinson, 2017; Clark, 1996; Clark & Schaefer, 1989; Holler & Levinson, 2019)). It is possible that language regions would respond differently when comprehending language spoken to the participant with the expectation of their response (e.g., a conversation with one other person), compared to a context in which they are observing others speaking and could not respond (see (Bögels & Levinson, 2017) for an excellent review of neuroimaging studies related to conversation).

Despite these limitations, the current task could have key applications. The original motivation for the stimulus section was to create a language task for use with young children. Extensive prior literature has demonstrated the benefits of naturalistic movie-based stimuli for young children (Cantlon, 2020; Cantlon & Li, 2013; Kamps et al., 2022; Redcay & Moraczewski, 2020; Richardson et al., 2018; Vanderwal et al., 2015, 2019). Given that this child-friendly task was engaging and effective for adults, it may be used across a wide age range and for other populations who may find classic language tasks hard to tolerate. In ongoing work, we are using this task to localize language processing in very young children and also to determine whether language regions are likewise insensitive to the differences between child-directed (monologue) and observed (dialogue) speech during language learning.

## Conclusion

Our results suggest that canonical left-hemisphere cortical language regions are not sensitive to features of multimodal dialogue other than comprehensibility of the speech. Processing non-linguistic features of dialogue is instead carried out by other brain regions and networks.

## Materials

Stimuli transcriptions and descriptions, analysis code, stimulus presentation code, processed data, and link to raw data on OpenNeuro can be found on OSF: https://osf.io/whsb7/. Raw stimuli can be provided upon request.

## Acknowledgements

We would like to thank Somaia Saba, Hana Ro, and Michelle Hung for their assistance with stimuli creation. Thank you to Ev Fedorenko, Shari Liu, and Nancy Kanwisher for helpful feedback on this manuscript. Thank you to Steve Shannon and Atsushi Takahashi at the Athinoula A. Martinos Imaging Center at MIT. Finally, thank you to our participants for making this research possible.

