## Supplemental Information for "Left-hemisphere cortical language regions respond equally to observed dialogue and monologue"

### Supplementary Materials

Additional Methods ... 1
Stimuli Characterization ... 1
fMRIprep specifications ... 2
Task performance ... 5
Deviations from Preregistration ... 5
Supplemental Figures ... 6
Supplementary Figure 1: SS-BlockedLang stimuli characteristics ... 6
Supplementary Figure 2: Whole brain contrast for social interaction ... 6
Supplementary Figure 3: SS-IntDialog average magnitude by condition within right language homologues ... 7
Supplemental Tables ... 7
Supplementary Table 1: SS-BlockedLang statistics in language regions including both hemispheres ... 7
Supplementary Table 2: SS-BlockedLang comprehensible dialogue regions ... 8
Supplementary Table 3: SS-IntDialog magnitude in language regions and right language homologues ... 9

#### Additional Methods

##### ***Stimuli Characterization***

To characterize the final stimuli, we transcribed each video and analyzed key linguistic features. We focused on stimuli features in Experimental Task 1 (SS-BlockedLang), as we wanted to identify potential differences in linguistic features between the monologue and dialogue blocks (a "block" consisted of either one 20-second video of dialogue, or two 10-second videos of monologue). Each block's audio was first transcribed by a computer-generated transcription service, then edited by a human for typos, punctuation, and removing other sounds from the transcript (e.g., stuttering and sound effects). Reading ease was calculated using the Flesch reading ease formula, which takes into account words per sentence and syllables per word, resulting in a value between 0 to 100, with 100 being the easiest to read (Flesch, 1948). Mean length of utterance (MLU) was calculated as the average number of words per sentence for each block, and Words Per Clip was the total number of words per block. The Flesch-Kincaid Reading Ease Score, MLU, and Words Per Clip were calculated using the following online calculator: https://goodcalculators.com/flesch-kincaid-calculator/. Conversational turns were counted based on the number of times characters in the dialogue blocks switched speaking (e.g., Abby-Elmo-Abby-Elmo = 3). We considered utterances to include vocalizations like "mm-hm" and "yeah" in addition to full sentences. Mental state terms were identified by an expert in this domain as terms that expressed mental states and emotions; we counted the total number of mental state terms per block. Average age of acquisition (AoA) corresponds to the age (in years) that a human learns a word, averaged across all the words per block (we used the "AoA Test Based score" from Brysbaert & Biemiller, 2017; see below for additional details). Transcriptions and values for all of these features are available on OSF: https://osf.io/whsb7/.

Additional details for calculating average AoA: Proper nouns (e.g., "Elmo") were not included. If a verb in a particular tense was not in the dictionary, it was assigned the score to the closest tense (e.g., "wants" was the word and "wants" was not in the dictionary but "want" was, so it was assigned the score for "want"). If a singular version of a noun was in the dictionary, the plural version in the transcript was counted as the singular rating. A human went through each clip to ensure that the correct score was assigned to each word in the block based on meaning, as some words had multiple possible definitions with different AoA scores. We removed words from the average calculation if they were not in the AoA dictionary (e.g., "yummy," "I'm," or "whatcha"). Helping verbs were collapsed into one unit and evaluated based on the main verb ("to" "help" became "to help"). The average score per block included scores for words as many times as they appeared in the block, including if the same word appeared in a block multiple times.

***fMRIprep specifications***

**Note that the following boilerplate description is copied verbatim from the fMRIprep output, per guidelines from the developers.*

Results included in this manuscript come from preprocessing performed using fMRIPprep 1.2.6 (Esteban, Markiewicz, et al. (2018); Esteban, Blair, et al. (2018); RRID:SCR_016216), which is based on Nipype 1.1.7 (Gorgolewski et al. (2011); Gorgolewski et al. (2018); RRID:SCR_002502).

**Anatomical data preprocessing**

The T1-weighted (T1w) image was corrected for intensity non-uniformity (INU) using N4BiasFieldCorrection (Tustison et al. 2010, ANTs 2.2.0), and used as T1w-reference throughout the workflow. The T1w-reference was then skull-stripped using antsBrainExtraction.sh (ANTs 2.2.0), using OASIS as target template. Brain surfaces were reconstructed using recon-all (FreeSurfer 6.0.1, RRID:SCR_001847, Dale, Fischl, and Sereno 1999), and the brain mask estimated previously was refined with a custom variation of the method to reconcile ANTs-derived and FreeSurfer-derived segmentations of the cortical gray-matter of Mindboggle (RRID:SCR_002438, Klein et al. 2017). Spatial normalization to the ICBM 152 Nonlinear Asymmetrical template version 2009c (Fonov et al. 2009, RRID:SCR_008796) was performed through nonlinear registration with antsRegistration (ANTs 2.2.0, RRID:SCR_004757, Avants et al. 2008), using brain-extracted versions of both T1w volume and template. Brain tissue segmentation of cerebrospinal fluid (CSF), white-matter (WM) and gray-matter (GM) was performed on the brain-extracted T1w using fast (FSL 5.0.9, RRID:SCR_002823, Zhang, Brady, and Smith 2001).

**Functional data preprocessing**

For each of the 10 BOLD runs found per subject (across all tasks and sessions), the following preprocessing was performed. First, a reference volume and its skull-stripped version were generated using a custom methodology of fMRIPrep. The BOLD reference was then co-registered to the T1w reference using bbregister(FreeSurfer) which implements boundary-based registration (Greve and Fischl 2009). Co-registration was configured with nine degrees of freedom to account for distortions remaining in the BOLD reference. Head-motion parameters with respect to the BOLD reference (transformation matrices, and six corresponding rotation and translation parameters) are estimated before any spatiotemporal filtering using mcflirt (FSL 5.0.9, Jenkinson et al. 2002). The BOLD time-series, were resampled to surfaces on the following spaces: fsaverage5. The BOLD time-series (including slice-timing correction when applied) were resampled onto their original, native space by applying a single, composite transform to correct for head-motion and susceptibility distortions. These resampled BOLD time-series will be referred to as preprocessed BOLD in original space, or just preprocessed BOLD. First, a reference volume and its skull-stripped version were generated using a custom methodology of fMRIPrep. Automatic removal of motion artifacts using independent component analysis (ICA-AROMA, Pruim et al. 2015) was performed on the preprocessed BOLD on MNI space time-series after removal of non-steady state volumes and spatial smoothing with an isotropic, Gaussian kernel of 6mm FWHM (full-width half-maximum). Corresponding "non-aggresively" denoised runs were produced after such smoothing. Additionally, the "aggressive" noise-regressors were collected and placed in the corresponding confounds file. The BOLD time-series were resampled to MNI152NLin2009cAsym standard space, generating a preprocessed BOLD run in MNI152NLin2009cAsym space. First, a reference volume and its skull-stripped version were generated using a custom methodology of fMRIPrep. Several confounding time-series were calculated based on the preprocessed BOLD: framewise displacement (FD), DVARS and three region-wise global signals. FD and DVARS are calculated for each functional run, both using their implementations in Nipype (following the definitions by Power et al. 2014). The three global signals are extracted within the CSF, the WM, and the whole-brain masks. Additionally, a set of physiological regressors were extracted to allow for component-based noise correction (CompCor, Behzadi et al. 2007). Principal components are estimated after high-pass filtering the preprocessed BOLD time-series (using a discrete cosine filter with 128s cut-off) for the two CompCor variants: temporal (tCompCor) and anatomical (aCompCor). Six tCompCor components are then calculated from the top 5% variable voxels within a mask covering the subcortical regions. This subcortical mask is obtained by heavily eroding the brain mask, which ensures it does not include cortical GM regions. For aCompCor, six components are calculated within the intersection of the aforementioned mask and the union of CSF and WM masks calculated in T1w space, after their projection to the native space of each functional run (using the inverse BOLD-to-T1w transformation). The head-motion estimates calculated in the correction step were also placed within the corresponding confounds file. All resamplings can be performed with a single interpolation step by composing all the pertinent transformations (i.e. head-motion transform matrices, susceptibility distortion correction when available, and co-registrations to anatomical and template spaces). Gridded (volumetric) resamplings were performed using antsApplyTransforms (ANTs), configured with Lanczos interpolation to minimize the smoothing effects of other kernels (Lanczos 1964). Non-gridded (surface) resamplings were performed using mri_vol2surf(FreeSurfer).

Many internal operations of fMRIPrep use Nilearn 0.5.0 (Abraham et al. 2014, RRID:SCR_001362), mostly within the functional processing workflow. For more details of the pipeline, see the section corresponding to workflows in fMRIPrep's documentation.

#### ***Task performance***

For SS-BlockedLang and SS-IntDialog, we recorded button presses after each video when a still image of Elmo appeared on the screen. Accuracy on the attention checks was 99.4% overall for SS-BlockedLang and 99.2% overall for SS-IntDialog. We preregistered that we would exclude runs if the participant missed more than 50% of the attention checks; no runs were excluded based on this criterion. For the ToM localizer, we measured accuracy on the true/false questions; accuracy was 77.5% overall. We preregistered that we would exclude runs with less than 70% accuracy on the ToM localizer. However, upon examining the accuracy after data collection, we realized that this would result in the exclusion of 1 run of ToM localizer from 10 of our 20 participants. Upon inspection, many participants with lower accuracy had no responses recorded for some trials, presumably because they responded to the true/false prompt after the recording window. Because this task was being used as a localizer and we had reason to believe that participants were aiming to read the blocks and answer the questions, we decided not to exclude trials based on accuracy.

#### ***Deviations from Preregistration***

(1) We used a different version of fMRIprep than specified in the preregistration. (2) We decided not to exclude runs of the ToM localizer task based on performance on the true/false questions. (3) We did not preregister testing for effects at the network level, but decided to include these tests along with effects at the ROI level. (4) Not all of the analyses specified in the preregistrations are included in this paper, including: analyses involving overlap or laterality index, analyses varying the significance threshold (we used our primary preregistered threshold of $p<.001$), analyses directly comparing the effect of matched vs. mismatched audio in SS-IntDialog in language vs. ToM regions, whole-brain analyses of forward monologue-specific effects, and analyses comparing variance explained in language regions’ response using the experimenter-derived regressor of forward/backward speech vs. average timecourse of within-group subjects in the SS-IntDialog task.

#### Supplemental Figures

***Supplementary Figure 1: SS-BlockedLang stimuli characteristics***

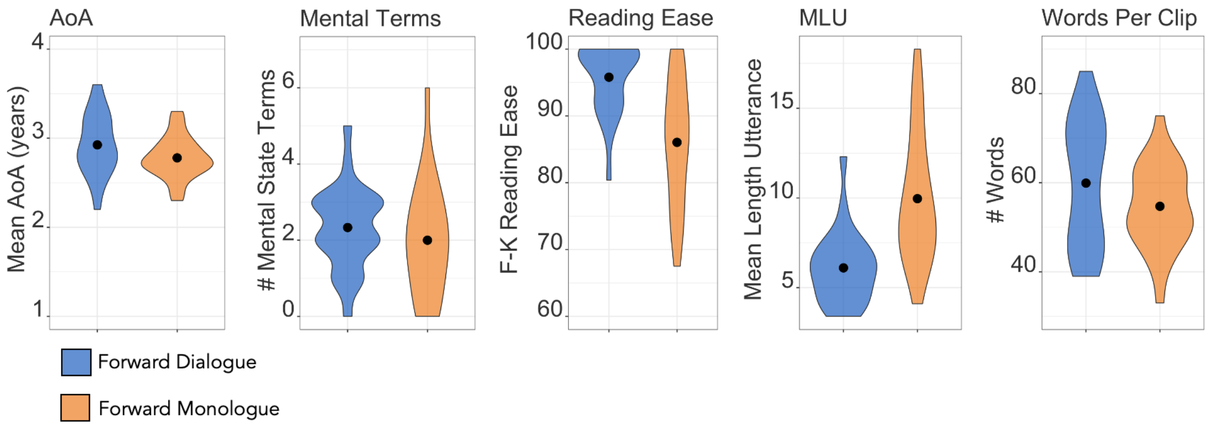


*Using Welch's two sample t-tests, there was no difference between dialogue and monologue for mental state terms (t-value=.89, p=.38), average age of acquisition for words in each block (t-value=1.72, p-value = 0.09), or total number of words per block (t-value=1.5, p=.14). Flesch-Kincaid reading ease score was higher for dialogue than monologue (t-value=4.48, p<.001), and monologues had more words per sentence (t-value=-4.56, p<.001).*

***Supplementary Figure 2: Whole brain contrast for social interaction***

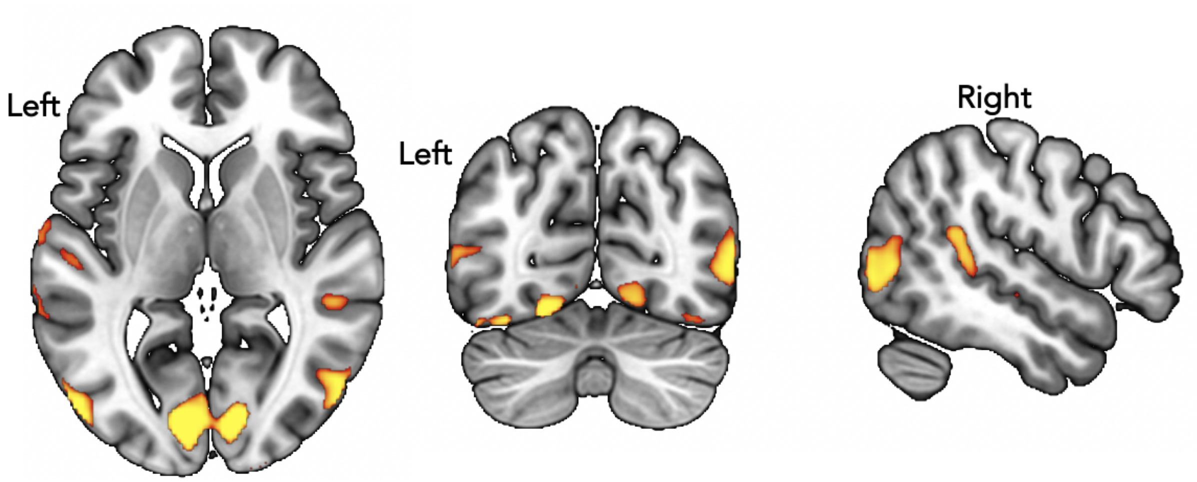


*Group whole-brain analysis for the Backward Dialogue > Backward Monologue contrast in SS-BlockedLang. Shown at threshold: p<.01, TFCE corrected.*

***Supplementary Figure 3: SS-IntDialog average magnitude by condition within right language homologues***

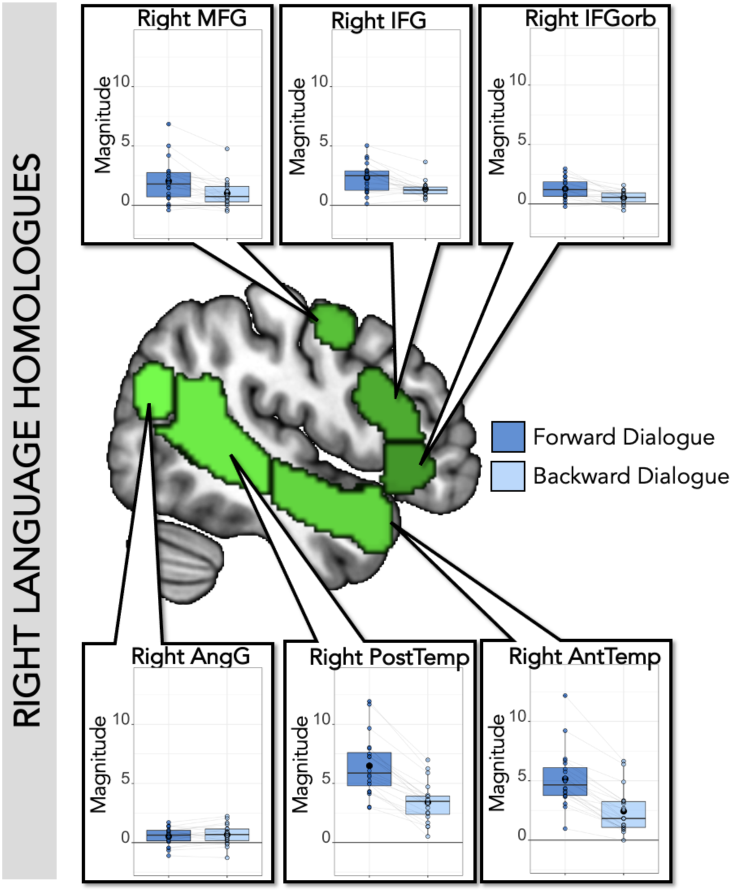


***Center***: *Right hemisphere language parcels overlaid on template brain (green; parcels include right IFGorb, IFG, MFG, AntTemp, PostTemp, and AngG from https://evlab.mit.edu/funcloc/).* ***Panels***: *Average response magnitude per individual for each condition in the SS-IntDialog task was extracted from subject-specific functional regions of interest for language (blue: Forward Dialogue; light blue: Backward Dialogue). All regions responded more to Forward than Backward speech. Each individual's datapoints are connected by light gray lines.*

#### Supplemental Tables

***Supplementary Table 1: SS-BlockedLang statistics in language regions including both hemispheres***

| ROI | Significant main effects and interactions |
|---|---|
| IFGorb | **FvB**: (Est.=0.57, S.E.=.05, t-value=11.72, p-value<.001);<br>**LvR**: (Est.=0.25, S.E.=0.05, t-value=5.24, p-value<.001);<br>**FvB*LvR**: (Est.=0.18, S.E.=0.05, t-value=3.65, p-value<.001) |

| | |
|---|---|
| IFG | **FvB**: (Est.=0.70, S.E.=0.06, t-value=12.40, p-value<.001);<br>**FvB*LvR**: (Est.=0.22, S.E.=0.06, t-value=3.95, p-value<.001) |
| MFG | **FvB**: (Est.=0.66, S.E.=0.07, t-value=9.80, p-value<.001)<br>**LvR**: (Est.=-0.21, S.E.=0.07, t-value=-3.13, p-value=0.002)<br>**FvB*LvR**: (Est.=0.24, S.E.=3.55, t-value=, p-value<.001) |
| AntTemp | **FvB**: (Est.=1.31, S.E.=0.08, t-value=15.51, p-value<.001)<br>**LvR**: (Est.=-0.53, S.E.=0.08, t-value=-6.30, p-value<.001) |
| PostTemp | **FvB**: (Est.=1.71, S.E.=0.13, t-value=13.56, p-value<.001) |
| AngG | **FvB**: (Est.=0.29, S.E.=0.07, t-value=3.84, p-value<.001) |

*lmer(mean_topvoxels_extracted~f_or_b*m_or_d*l_or_r+(1|participantID), REML = FALSE). Uncorrected p-value. Significant at level p<.05 when Bonferroni corrected for 6 ROIs (p<.0083). Only shows significant main effects and interactions per ROI.*

***Supplementary Table 2: SS-BlockedLang comprehensible dialogue regions***

| Exploratory: Comprehensible Dialogue Regions | | | | | | | |
|---|---|---|---|---|---|---|---|
| Significant Clusters | | | | | Results from independent ss-fROIs | | |
| **ROI** | **Voxels** | **MAX T-value** | **Peak X,Y,Z (mm)** | **COG X,Y,Z (mm)** | **Forward v. Backward** | **Dialogue v. Monologue** | **Interaction** |
| Right Temporal Pole | 299 | 6.64 | 54, 18, -26 | 52, 12.6, -30.9 | Est.=0.49; S.E.=0.05; t-value=10.40 | Est.=0.23; S.E.=0.05; t-value=4.82 | Est.=0.16; S.E.=0.05; t-value=3.41 |
| Right STS | 98 | 6.33 | 50, -26, -6 | 52, -23.9, -6.18 | Est.=0.68; S.E.=0.09; t-value=7.83 | Est.=0.28; S.E.=0.09; t-value=3.21 | Est.=0.16; S.E.=0.09; t-value=1.79 |
| Left Crus II (Cerebellum) | 29 | 4.61 | -26, -78, -34 | -28.1, -80, -34.6 | Est.=0.21; S.E.=0.05; t-value=4.67 | Est.=0.11; S.E.=0.05; t-value=2.43 | Est.=0.12; S.E.=0.05; t-value=2.65 |
| Left STS | 14 | 4.74 | -50, -30, -4 | -50.3, -30.6, -3.42 | Est.=0.72; S.E.=0.06; t-value=11.92 | Est.=0.16; S.E.=0.06; t-value=2.58 | Est.=0.12; S.E.=0.06; t-value=1.92 |

*Significant clusters at p<.001 (uncorrected, df=19, two-tailed). Peak coordinates and center of gravity (COG) for the cluster (weighted average of the coordinates by the intensities within the cluster). No significant voxels at p<.001 TFCE corrected. Within ss-fROIs defined based on 10mm spheres around the group cluster COG coordinates, there was a higher response to*

*Forward than Backward speech in all regions, a higher response to Dialogue speech compared to Monologue speech in all regions, and an interaction in Right Temporal Pole and Left Crus II (cerebellum). Results (Est. = estimate, S.E. = standard error, t-value) from the model: lmer(mean_topvoxels_extracted~f_or_b*d_or_m+(1|participantID), REML = FALSE). P-values not reported since analyses were exploratory.*

***<u>Supplementary Table 3: SS-IntDialog magnitude in language regions and right language homologues</u>***

| ROI | Forward | Backward | Forward>Backward |
|---|---|---|---|
| Left IFGorb | M(SD) = 2.68(1.43);<br>range = 0.54-6.14 | M(SD) = 0.73(0.60);<br>range = -0.17-2.06 | Est.= 0.98<br>S.E.= 0.12<br>t-value= 8.34<br>**p-value<.001 *** |
| Left IFG | M(SD) = 3.63(1.10);<br>range = 1.89-5.89 | M(SD) = 1.30(0.64);<br>range = 0.39-2.38 | Est.= 1.16<br>S.E.= 0.12<br>t-value= 9.71<br>**p-value<.001 *** |
| Left MFG | M(SD) = 3.08(1.21);<br>range = 1.35-6.52 | M(SD) = 0.38(0.68);<br>range = -0.70-1.87 | Est.= 1.35<br>S.E.= 0.11<br>t-value= 12.70<br>**p-value<.001 *** |
| Left AntTemp | M(SD) = 4.36(1.61);<br>range = 2.49-9.00 | M(SD) = 1.45(1.10);<br>range = 0.04-4.68 | Est.= 1.45<br>S.E.= 0.10<br>t-value= 14.20<br>**p-value<.001 *** |
| Left PostTemp | M(SD) = 7.48(2.32);<br>range = 4.65-13.84 | M(SD) = 2.99(1.69);<br>range = 0.70-7.43 | Est.= 2.24<br>S.E.= 0.11<br>t-value= 21.29<br>**p-value<.001 *** |
| Left AngG | M(SD) = 1.37(1.44);<br>range = -1.05-4.45 | M(SD) = 0.50(1.12);<br>range = -1.21-4.10 | Est.= 0.43<br>S.E.= 0.09<br>t-value= 5.07<br>**p-value<.001 *** |
| Right IFGorb | M(SD) = 1.26(0.88);<br>range = -0.22-2.96 | M(SD) = 0.52(0.59);<br>range = -0.55-1.58 | Est.= 0.37<br>S.E.= 0.09<br>t-value= 4.30<br>**p-value<.001 *** |
| Right IFG | M(SD) = 2.32(1.26); | M(SD) = 1.35(0.68); | Est.= 0.49 |

| | | | |
|---|---|---|---|
| | range = 0.11-5.03 | range = 0.45-3.65 | S.E.= 0.12<br>t-value= 4.15<br>**p-value<.001 *** |
| Right MFG | M(SD) = 2.04(1.79);<br>range = -0.41-6.85 | M(SD) = 1.01(1.16);<br>range = -0.49-4.76 | Est.= 0.51<br>S.E.= 0.13<br>t-value= 3.98<br>**p-value<.001 *** |
| Right AntTemp | M(SD) = 5.15(2.42);<br>range = 0.97-12.17 | M(SD) = 2.44(1.83);<br>range = -0.01-6.65 | Est.= 1.36<br>S.E.= 0.11<br>t-value= 11.87<br>**p-value<.001 *** |
| Right PostTemp | M(SD) = 6.49(2.53);<br>range = 2.96-11.94 | M(SD) = 3.41(1.67);<br>range = 0.52-6.99 | Est.= 1.54<br>S.E.= 0.15<br>t-value= 10.57<br>**p-value<.001 *** |
| Right AngG | M(SD) = 0.53(0.76);<br>range = -1.10-1.71 | M(SD) = 0.67(0.85);<br>range = -1.28-2.24 | Est.= -0.07<br>S.E.= 0.05<br>t-value= -1.34<br>p-value=0.19 |

*Average magnitude (betas) for SS-IntDialog conditions (Forward and Backward), extracted from ss-fROIs for language based on the auditory language localizer. Results (Est. = estimate, S.E. = standard error, t-value, and uncorrected p-value) from the model: lmer(mean_topvoxels_extracted~f_or_b +(1|participantID), REML = FALSE).*
** indicates significance level p<.05, Bonferroni corrected for 6 ROIs (p<.0083)*
